# Mutant GNAS drives a pyloric metaplasia with tumor suppressive glycans in intraductal papillary mucinous neoplasia

**DOI:** 10.1101/2024.02.25.581948

**Authors:** Vincent Quoc-Huy Trinh, Katherine E. Ankenbauer, Sabrina M. Torbit, Jiayue Liu, Maelle Batardiere, Bhoj Kumar, H. Carlo Maurer, Frank Revetta, Zhengyi Chen, Angela Kruse, Audra Judd, Celina Copeland, Jahg Wong, Olivia Ben-Levy, Brenda Jarvis, Monica Brown, Jeffrey W. Brown, Koushik Das, Yuki Makino, Jeffrey M. Spraggins, Ken S. Lau, Parastoo Azadi, Anirban Maitra, Marcus C.B. Tan, Kathleen E. DelGiorno

**Author notes:** Correspondence to: KE DelGiorno, Department of Cell and Developmental Biology, Vanderbilt University. These authors contributed equally (Vincent Quoc-Huy Trinh, Katherine E. Ankenbauer).

## Abstract

**BACKGROUND & AIMS:** Intraductal Papillary Mucinous Neoplasms (IPMNs) are cystic lesions and *bona fide* precursors for pancreatic ductal adenocarcinoma (PDAC). Recent studies have shown that pancreatic precancer is characterized by a transcriptomic program similar to gastric metaplasia. The aims of this study were to assay IPMN for pyloric markers, to identify molecular drivers, and to determine a functional role for this program in the pancreas.

**METHODS:** Pyloric marker expression was evaluated by RNA-seq and multiplex immunostaining in patient samples. Cell lines and organoids expressing *Kras^G12D^* +/- *GNAS^R201C^* underwent RNA sequencing. A PyScenic-based regulon analysis was performed to identify molecular drivers, and candidates were evaluated by RNA-seq, immunostaining, and small interfering RNA knockdown. Glycosylation profiling was performed to identify *GNAS^R201C^*-driven changes. Glycan abundance was evaluated in patient samples.

**RESULTS:** Pyloric markers were identified in human IPMN. *GNAS^R201C^*drove expression of this program as well as an indolent phenotype characterized by distinct glycosyltransferase changes. Glycan profiling identified an increase in LacdiNAcs and loss of pro-tumorigenic Lewis antigens. Knockdown of transcription factors *Spdef* or *Creb3l1* or chitinase treatment reduced LacdiNAc deposition and reversed the indolent phenotype. LacdiNAc and 3’-sulfoLe^A/C^ abundance discriminated low from high grade patient IPMN.

**CONCLUSION:** *GNAS^R201C^* drives an indolent phenotype in IPMN by amplifying a differentiated, pyloric phenotype through SPDEF/CREB3L1 which is characterized by distinct glycans. Acting as a glycan rheostat, mutant *GNAS* elevates LacdiNAcs at the expense of pro-tumorigenic acidic Lewis epitopes, inhibiting cancer cell invasion and disease progression. LacdiNAc and 3’-Sulfo-Le^A/C^ are mutually exclusive and may serve as markers of disease progression.

## INTRODUCTION

Pancreatic ductal adenocarcinoma (PDAC) is currently the third leading cause of cancer-related deaths in the United States and is slated to become second by the year 2040^1^. This is largely due to late detection – pancreatic tumors grow silently, approximately 85% of patients present with incurable locally advanced or metastatic disease, and distant metastatic spread can occur when the tumors are small (<5mm). On the other hand, the time required for PDAC to emerge from a normal cell is long, some 15-20 years^2^. Thus, improving our understanding of the early events in neoplastic transformation is necessary and crucial to allow earlier diagnosis of PDAC, and also important therapeutically, to gain insights into how progression to cancer can be blocked or even reversed.

A major obstacle to the study of early events in pancreatic carcinogenesis is that the main precursor to PDAC, pancreatic intraepithelial neoplasia (PanIN), is clinically silent and radiographically occult. Intriguingly, the opposite is true of intraductal papillary mucinous neoplasms (IPMN), which contribute to approximately 25% of cases of PDAC^3^. As IPMN are cystic rather than microscopic like PanIN, they are easily identified on abdominal imaging scans and as a result, 90% are diagnosed before cancer is present. Thus, for most IPMN, there is a window of opportunity for surveillance and intervention before invasive disease develops^4, 5^. For all pre-malignant tumors, accurate assessment of the risk of malignant transformation is essential for clinical decision-making. This is particularly true of IPMN, because the risks of both under-and over-treatment are high. If cancer risk is under-estimated and the patient is surveilled, then the emergence of PDAC could be lethal. Conversely, surgery for IPMN carries with it very significant risks – the post-operative complication rate is 30-50% and the 90-day mortality is 3-5%^6–8^. Unfortunately, markers distinguishing low and high grade IPMN are currently lacking, resulting in an abundance of unnecessary surgeries. Approximately 50% of patients undergoing surgery for IPMN are found to lack PDAC upon final pathology. Therefore, better markers are desperately needed for the risk stratification of IPMN patients.

Oncogenic mutations in *KRAS* are common in human PanIN lesions and have been shown to drive tumorigenesis through increasingly dysplastic grades of PanIN to PDAC in genetically engineered mouse models (GEMMs)^9^. Like PanIN, IPMN harbor *KRAS* mutations (∼80%) but, distinct from PanIN, approximately two-thirds also express oncogenic GNAS (∼66%) driver mutations^10–12^. In GEMMs, combined expression of both mutant *KRAS* and *GNAS* has been shown to drive PDAC formation through a mixed phenotype of PanIN and IPMN^13^. In cell lines, *GNAS^R201H/C^* mutations have been shown to drive an indolent, less aggressive phenotype, potentially by limiting *KRAS* signaling^13, 14^. Recent studies, however, have shown that high grade human IPMN bearing *GNAS* mutations are accompanied by a loss of the wild type allele, suggesting that allelic imbalance drives disease. Further, these IPMN are accompanied by additional mutations driving MAPK signaling and tumor suppressor loss; *GNAS* mutations alone are likely insufficient to drive IPMN progression^15, 16^.

Metaplasia is considered to be an initiating event in both PanIN and IPMN formation. Metaplasia is a pathological term for the transdifferentiation of one cell type to another and is a form of plasticity common to injury and oncogene-induced disease progression in the gastrointestinal tract^17–19^. While it is largely thought to represent a reactive tissue response to mitigate injury, it is also considered to be the first step in tumorigenesis in several organs. Metaplasia falling under the general rubric of pyloric-like has been reported in stomach injury and tumorigenesis, serrated colorectal polyps, and pancreatic injury and tumorigenesis occurring through PanIN progression^20–24^. Metaplasia in the stomach, or spasmolytic polypeptide expressing metaplasia (SPEM), is the best characterized and reflects the transition of gastric chief cells (digestive enzyme producing cells) to a phenotype described by the expression of specific markers (e.g. MUC6, TFF2, AQP5, CD44v9, GKN3, etc.)^25–27^. When SPEM is accompanied by the presence of a foveolar pit cell lineage (TFF1, GKN1, GKN2, MUC5AC), the phenotype is a pyloric metaplasia due to a recapitulation of the pylorus region of the stomach^28, 29^. Recently, we combined single cell RNA-sequencing (scRNA-seq), electron microscopy, and histopathology and showed that injury-induced acinar to ductal metaplasia (ADM) in the pancreas results in the formation of chemosensory tuft cells, hormone-producing enteroendocrine cells, and a population bearing canonical markers of SPEM^20^. ADM and PanIN resulting from oncogenic *Kras^G12D^* expression are also characterized by SPEM with the additional formation of a separate, distinct foveolar pit cell-like population, reflecting pyloric metaplasia^20, 21^. The functional role of individual pyloric markers has previously been studied in GEMMs of gastric disease^30, 31^; however, the program itself is hypothesized to represent a shared mechanism by which gastrointestinal organs respond to injury and tumorigenesis^29^.

Here, we assayed models of IPMN and patient samples for the expression of pyloric markers. We show that mutant *GNAS* is sufficient to amplify a pyloric phenotype, including SPEM and pit cell markers, mucus production, and distinct glycosylation changes, and identify transcription factors SPDEF and CREB3L1 as critical regulators of this program. Finally, we show that *GNAS^R201C^*-induced glycan changes contribute to the indolent phenotype and may be used to distinguish high from low grade IPMN to risk stratify IPMN patients.

## MATERIALS AND METHODS

### Human Samples

A cohort of 41 patients with intraductal papillary mucinous neoplasms (IPMNs) ranging from low-grade dysplasia, high-grade dysplasia, to invasive were selected from Vanderbilt University Medical Center’s institutional cohort with institutional review board approval (#101066). Normal stomach body, small intestine, and colon tissues were used as controls to threshold signal intensity.

### Mice

Mice were housed in accordance with National Institutes of Health guidelines in American Association for Accreditation of Laboratory Animal Care-accredited facilities at Vanderbilt University or M.D. Anderson Cancer Center. The Vanderbilt or M.D. Anderson Institutional Animal Care and Use Committees (IACUC) approved all animal studies. *LSL-Kras^G12D/+^*, *Ptf1a^Cre/+^*, *LSL-rtTA-TetO-GNAS^R201C^*(*Kras;GNAS*) mice have previously been described^9, 13, 32^. Mice were given a diet containing doxycycline (Tusculum Feed Center, #9205-0827) beginning at 8 weeks of age for a period of either 10 or 20 weeks.

### Cell culture and *Gnas^R201C^* induction

Murine cell lines generated from *Kras;GNAS* mice^13^ (4838 and C241) were cultured in RPMI with L-glutamine (Corning 10-040-CV), 10% Tet system approved FBS (Gibco A4736401) and 1 x antibiotic/antimitotic (Corning 30-004-CI). To induce human *GNAS^R201C^* expression, cells were stimulated with 1 μg/ml doxycycline (RPI Research Products International D43020) for 48-72 hours. *Spdef* or *Creb3l1* knockdown was accomplished by siRNA mediated knockdown. siRNA for *Spdef* (#184152, Thermo-Fisher), *Creb3l1* (#162495, Thermo-Fisher), or a nonspecific control (*Silencer* Negative Control No. 2 siRNA, Thermo-Fisher) were transfected using lipofectamine RNAiMAX (#13778075, Thermo-Fisher). Cells were seeded in 6cm dishes with 1μg/ml of doxycycline for 48 hours. At 40% confluency, cells were transfected for a final concentration of 20 pmol. After 72 hours, RNA was extracted, or cells were prepared for immunofluorescence. Chitinase treatment was performed in a similar matter with sells being seeded in 6-well plates with 1μg/ml of doxycycline for 48 hours. At 60% confluency, cells were treated with chitinase from *streptomyces griseus* (Sigma-Aldrich, C6137). After 48 hours, RNA was extracted, and cells were prepared for immunofluorescence. Western blotting was conducted using Bio-Rad Mini-Protean 4-20% TGX gels, PVDF membranes and the Trans Blot Turbo system. Blocking was done in 5% non-fat milk. Primary antibodies are listed in **Table S1**. Blots were imaged using Immobilon HRP Substrate (WBKLS0500) and processed using an Amersham AI600 imaging system. For quantitative real-time polymerase chain reaction (qRT-PCR), RNA was isolated using Quick-RNA MiniPrep kits (Zymo Research R1055), and concentration was measured using NanoDrop 2000. qRT-PCR was performed using a Luna Universal One-Step RT-qPCR Kit (New England Laboratories, E3005) and analyzed using Bio-Rad CFX96 system and CFX Manager 3.1. Primers are listed in **Table S2**. For immunofluorescence, cells were seeded on a Matrigel-coated coverslip (1:50 dilution; Corning, 356231) and fixed with 4% PFA (Electron Microscopy Services, 15712) for 15 minutes at room temperature. Following fixation, cells were washed 5 times in PBS and blocked and permeabilized using buffer containing 1% BSA (w/v), 5% donkey serum (v/v), 5% goat serum (v/v) and 0.3% Triton-X for 30 minutes at room temperature. Coverslips were washed in PBS 5 times. For primary incubation, primary antibodies (**Table S1**) were diluted in blocking buffer (5% donkey serum, 1% BSA and 0.05% Triton-X) and incubated at 37°C for 2 hours. Coverslips were washed 5 times in PBS and secondary antibody diluted in blocking buffer was added. Secondary antibodies were incubated at 37°C for 1 hour. Coverslips were washed 5 times in PBS, mounted with ProLong Gold Antifade Mountant with DAPI (Invitrogen, P36931) and imaged with a 20x objective on an Olympus VS200 Slide Scanner (Olympus, Tokyo, Japan). Invasion assays were performed using Corning BioCoat Matrigel Invasion Chambers with 8.0 µm PET Membrane (Corning, 354480). Assays were prepared according to manufacturer protocol and 125,000 cells were plated per well. After 24 hours, cells on the membrane were fixed with 4% PFA for 15 minutes and stained with 0.2% w/v crystal violet for 15 minutes. Membranes were carefully cut out of the inserts and mounted and imaged with a 20x objective on an Olympus VS200 Slide Scanner.

### Organoid generation, culture, and treatment

Organoids were established from the pancreata of 4, 6-week-old *Kras;GNAS* mice and 2, 6-month-old *GNAS* mice fed a normal chow diet. Briefly, pancreata were dissected, minced in Hank’s Balanced Salt Solution (HBSS) and incubated in digestion media containing 1 mg/mL collagenase IV (Gibco), 10 µg/mL of DNAse I, 0.5 mg/mL of soybean trypsin inhibitor (Gibco), and 5 mLs of dispase (StemCell) diluted in HBSS. Tissues were digested for 8-10 cycles by incubating in digestion media with gentle shaking at 37°C for 10-20 minutes per cycle. At the end of each cycle, cells were collected, the digestion buffer was neutralized with wash buffer (10% FBS in HBSS) and cells were placed on ice. After the last digestion, dispersed cells were filtered through a 70 <u>µm strainer and</u> pelleted at 1,000 x g for 5 minutes, embedded in Matrigel (Corning, cat# 356231) and organoid media was added. Pancreatic organoid media consisted of Advanced DMEM/F12 (Gibco) supplemented with 10% Noggin conditioned media, 10% R-spondin conditioned media, 1X B27 (Gibco), 10 mM nicotinamide (Sigma Aldrich), 1.25 mM N-acetylcysteine (Sigma Aldrich), 50 ng/mL of mEGF (Biolegend), 100 ng/mL of mFGF-10 (Biolegend), 10 nM hGastrin I (Anaspec), and 500 nM of A83-01 (Tocris). Y-27632 (Rock inhibitor, HelloBio) was added to media directly after isolation but removed in following passages. Organoids were maintained in 5% CO_2_ at 37°C and passaged every 4-7 days. To induce GNAS^R201C^ expression, organoids were treated with 2 µg/mL doxycycyline for 72 hours. RNA was extracted following manufacturer’s protocol (Zymo).

### Immunofluorescence analysis

Cell line IF images were analyzed for corrected total cell fluorescence (CTCF) using Fiji (National Institutes of Health)^33^. Briefly, representative ROIs in each image were selected using the freehand selection tool in the experimental channel. The area, mean, integrated density, and raw integrated density were calculated for each ROI. The same values were also taken from the background. To calculate the CTCF, the following formula was used: raw integrated density – (area*average background mean). For the area analysis, images were split into separate channels, thresholded, and the area of both the DAPI and experimental channel were measured. To normalize for the number of cells in each ROI, the area of the antibody or lectin stain was normalized to the area of the nuclear stain.

### Multiplex Immunohistochemistry (MxIHC)

Formalin-fixed paraffin-embedded tissues from the patient cohort were sectioned at 4 μm, heated to 60 °C for 30 min, deparaffinized in xylenes, and rehydrated in an ethanol gradient. Slides were stained with Mayer hematoxylin (Sigma-Aldrich, MHS32), coverslipped with 30% glycerol (Sigma-Aldrich, G5516) in phosphate-buffered saline (PBS) and scanned with an Olympus VS200 slide scanner (Olympus, Tokyo, Japan). Slides were de-coverslipped by immersion in PBS and underwent antigen retrieval in 10 mM sodium citrate pH 6.0 with a microwave set at maximum power until boiling bubbles appeared, reduced to minimum power for 20 minutes, and left at RT for 30 minutes. Slides were treated with 3% hydrogen peroxide for 10 minutes, blocked with Protein Block, Serum-Free (Agilent Dako, X090930-2) for 10 minutes, and incubated with the primary antibody overnight. Secondary antibodies were incubated for 1 hour, and the signal was revealed with AEC+ High Sensitivity Substrate Chromogen (Dako Agilent, K3469). The slides were then coverslipped with 30% glycerol (Sigma-Aldrich, G5516) in phosphate-buffered saline (PBS). Slides were scanned with the VS200. Slides were de-coverslipped by immersion in PBS and underwent a 2-minute double-distilled water, 2 minutes 70% ethanol, 2 minutes 95% ethanol, 2 minutes ethanol, 2-minute double-distilled water sequence to eliminate the 3-amino-9-ethylcarbazole (AEC) chromogen. Slides then restarted the previous sequence at the antigen retrieval step until all primary antibodies (**Table S1**) were performed. Scans were loaded in QuPath v0.4.0^34^ and registered with the image-combiner v0.3.0 package.

### Standard histological staining

Murine and patient tissues were cut in 5 µm or 4 µm sections, respectively, mounted, and stained as previously described^20^. Sections were deparaffinized in xylenes, rehydrated in a series of graded ethanols, and then washed in PBST and PBS. Endogenous peroxidase activity was blocked with a 1:50 solution of 30% H_2_O_2_:PBS followed by microwave antigen retrieval in 100 mM sodium citrate, pH 6.0. Sections were blocked with 1% bovine serum albumin and 5% normal goat serum in 10 mM Tris (pH 7.4), 100 mM MgCl_2_, and 0.5% Tween-20 for 1hr at room temperature. Primary antibodies (**Table S1**) were diluted in blocking solution and were incubated on tissue sections overnight. Slides were then washed, incubated in streptavidin-conjugated secondaries (Abcam), and developed with DAB substrate (Vector). Periodic Acid Schiff/Alcian Blue (PAS/AB) staining was performed per the manufacturer’s instructions (Abcam, ab245876). For standard IHC on human tissue sections, antigen retrieval was performed with pH 6.0 citrate buffer in a pressure cooker at 105°C for 15 minutes, with a 10-minute cool down. Blocking was performed in 0.03% H_2_O_2_ containing sodium azide for 5 minutes and primary antibodies (**Table S1**) were incubated for 60 minutes before detection (Dako EnVision+ System-HRP labeled Polymer) for 30 minutes and development for 5 minutes. All slides were scanned on the VS200.

### IHC quantification

<u>MxIHC Quantification.</u> Areas measuring 250 x 250 μm^2^ were acquired blindly on the hematoxylin layer, querying normal ducts, acinar-to-ductal metaplasia (ADM), low-grade dysplasia (LG), high-grade dysplasia (HG), and invasive disease under the supervision of a board-certified pathologist (VQT). Up to 3 areas of normal, ADM and INV, and up to 5 areas of LG and HG, were selected per patient (when present). An automated watershed threshold pixel detection was adjusted based on the staining patterns on the control gastric, small intestinal, and colonic tissues. Automated AEC deconvolution was performed in QuPath v0.4.0 by loading them as a H-DAB slide. An automatic calculation of the surface area with the AEC color deconvolution intensity over the threshold was then generated. Pseudocolors were given to each stain for the figures. <u>Monoplex Quantification of SPDEF, CREB3L1 and CREB3L4.</u> Scans of SPDEF, CREB3L1, and CREB3L4 were obtained with the VS200 and loaded in QuPath v0.4.3^34^. Color deconvolution of the hematoxylin stain was performed to select up to 3 areas of normal ducts, ADM, LG, HG, and INV. By consensus, one student and a board-certified pathologist (VQT) scored the nucleus and cytoplasm of cells on a scale of 0-1-2-3. <u>Cell type quantification in murine IPMN.</u> Scans of H&E staining from *Kras;GNAS* mice +/- doxycycline chow for 10 or 20 weeks (n = 3/condition, n = 12 total) were scored by a pathologist (VQT) and areas of IPMN and/or PanIN lesions were identified (n = 3, up to 6 regions/slide). Serial sections were stained for DCLK1 (tuft cells), synaptophysin (enteroendocrine cells), or PAS/AB (all mucins) and 1-6 lesions were identified per area of IPMN and/or PanIN (when present). All steps of analysis were performed blinded. Positive DCLK1 or synaptophysin cells were manually counted and divided by the number of nuclei per lesion to identify percent of that lesion constituted by a given cell type. To quantify PAS/AB mucin staining, lesions were annotated as regions of interest and manually drawn in FIJI. Total signal and PAS/AB signal were thresholded to include only stained areas and percent positive area was calculated.

### Glycosylation profiling of cell lines

Cells were lysed using 50 mM ammonium bicarbonate buffer, were treated with DTT, and then incubated at 50 °C for 30 min. The samples were then desalted by centrifugal filtration and ultrasonicated. BCA was used to quantify protein, and an equal amount of protein was further processed for N-glycan release. Samples were then treated with PNGaseF (37 °C for 48 h) and released N-glycans were collected using 10kDa cutoff spin filter, followed by lyophilization. O-glycoproteins were retrieved from the top of the spin filter and lyophilized before proceeding to β-elimination. Both released N- and O-glycans were permethylated and analyzed by ESI-MS (Orbitrap Eclipse Tribrid mass spectrometer). Glycan abundance (%) was calculated separately for each sample based on peak intensity. The N-glycan and O-glycan structures were assigned based on precursor masses (Sodiated) and mammalian biosynthetic pathway. Proposed structures are based on biosynthetic pathways and do not reflect MS/MS, compositional or linkage analysis.

### Imaging glycosylation mass spectrometry

FFPE tissue sections mounted on slides were incubated for one hour at 60°C followed by deparaffinization using a wash series consisting of: xylenes for 3 minutes, xylenes for 3 minutes, 100% ethanol for 1 minute, Carnoy’s solution (60% ethanol, 30% chloroform and 10% glacial acetic acid) for 3 minutes, Carnoy’s solution for 3 minutes, Carnoy’s solution for 3 minutes, 100% ethanol for 1 minute, 95% ethanol for 1 minute, 70% ethanol for 1 minute, 150 mM ammonium formate for 1 minute, 150 mM ammonium formate for 1 minute. Samples were placed in a vacuum desiccator for 10 minutes. Autofluorescence microscopy was collected from each slide using a Zeiss AxioScan Z1 slide scanner with a 10x objective using the DAPI (excitation wavelength 353 nm), GFP (excitation wavelength 488 nm), and dsRed (excitation wavelength 572 nm) filter sets using exposure times of 20 ms, 60 ms, and 250 ms, respectively. Thermal denaturation was performed in 10 mM citraconic anhydride for 30 minutes in a preheated steamer followed by buffer exchange with milli-q water. Samples were placed in a vacuum desiccator for 10 minutes. Samples were then coated with 1 µg/µL PNGase F Prime in HPLC-grade water using an HTX M5 robotic sprayer using the following conditions: 37°C nozzle temperature, ambient stage temperature, 10 psi-compressed air, 15 passes, 1200 mm/min velocity, 3 mm offset, CC spray pattern, 40 mm nozzle height, no dry time. A syringe pump equipped with a 500 µL Hamilton syringe with 3.26 mm diameter was used for enzyme spraying with a 25 µL/min flow rate. Enzymatic digestion was performed in a preheated digestion chamber containing 5 mL milli-q water at 38°C for 2 hours. α-Cyano-4- hydroxycinnamic acid matrix solution at a concentration of 7 mg/mL in 50% acetonitrile containing 0.1% TFA was applied to the slides using an HTX M5 sprayer with the following conditions: 72°C nozzle, 10 psi compressed air, ambient stage temperature, 0.1 mL/min flow rate, CC spray pattern, no dry time, 40 mm nozzle height, 10 passes, 1300 mm/min, 3 mm offset. All robotic spraying steps were performed at 76°F and 60% relative humidity. MALDI IMS was performed using a Bruker timsTOF Pro in positive ionization mode with an *m/z* from 500-4000, 200 laser shots at a frequency of 10,000Hz with a laser fluence of 39% and a pitch of 20 µm.

### Analysis of published single-cell RNA sequencing datasets

Processed count matrices from Bernard et al., were downloaded from the Gene Expression Omnibus (GEO) database^35^. Data analyses were executed using a dual-language approach, encompassing R (version 4.3.1, 2023-06-16) and Python (version 3.9.13, 2022-08-25), both tailored for a 64-bit macOS Ventura 13.4 platform. R integrated packages utilized in the study comprised ‘ggplot2 (3.4.3)’, ‘dplyr (1.1.3)’, ‘patchwork (1.1.3)’, ‘SeuratObject (4.1.3)’, ‘Seurat (4.3.0.1)’, ‘umap (0.2.10.0)’, ‘stxBrain.SeuratData (0.1.1)’, and ‘SeuratData (0.2.2)’. Python libraries of interest included ‘anndata (0.9.2)’, ‘scanpy (1.9.5)’, and ‘pandas (2.1.1)’. Seurat objects were established and normalized; individual Seurat objects representative of the six patient datasets from Bernard et al., were synthesized and post-merged into a unified object. Consolidated data were filtered to cells with a minimum of 300 genes, manifesting less than 15% mitochondrial genes (percent.mt<0.15). Emphasis was subsequently placed on identifying and leveraging highly variable features, with the selection of top 2,000 features (nFeature_RNA > 2000) via the ‘FindVariableFeatures()’ function. Post scaling, data was subjected to Principal Component Analysis (PCA) to demarcate inherent patterns. Exploiting the dimensionality reduced space (spanning the first 20 principal components), cellular neighborhoods were mapped with the ‘FindNeighbors()’ function (dims = 1:20,k.param = 20) and clusters were discerned using ‘FindClusters()’ with a resolution parameter set at 0.5. For downstream analysis, the MuDataSeurat tool, accessible on [GitHub] (https://github.com/PMBio/MuDataSeurat), was employed to convert Seurat objects into the ‘.h5ad’ format compatible with the ‘Scanpy’ library. UMAPs labeled by sample or for individual gene markers (‘CD44’,‘MUC2’,‘TFF2’,‘PTPRC’,‘EPCAM’,‘AQP5’,‘MUC5AC’,etc.) were plotted for visualization.

### Bulk human RNA sequencing analysis

Compartment-specific gene expression profiles of human IPMN (n = 19), PanIN (n = 26) and PDAC (n = 197) were generated using laser capture microdissection with subsequent RNA sequencing as previously described^36–38^. Differentially expressed genes between human IPMN and PanIN were identified leveraging a generalized linear model as implemented in DESeq2 R package^39^ and genes exhibiting a false discovery rate ≤ 0.05 were considered significantly differentially expressed.

### Bulk murine cell line RNA collection and sequencing analysis

RNA was extracted from either 4838 or C241 cells (+/- doxycycline) or organoid lines using Quick-RNA MiniPrep (Zymo Research R1055) and quality was measured using an Agilent Bioanalyzer; RNA Qubit assay was performed to measure RNA quantity. Poly(A) RNA enrichment was conducted using NEBNext Poly(A) mRNA Magnetic Isolation Module (NEB E7490), and the sequencing library was constructed using the NEBNext Ultra II RNA Library Prep Kit (E7765L) following the manufacturer’s instructions. End repair, A-tailing, and adapter ligation was performed to generate the final cDNA library. Library quality was assessed using a Bioanalyzer and quantified using a qPCR-based method with the KAPA Library Quantification Kit (KK4873) and the QuantStudio 12K instrument. 150 bp paired-end sequencing was performed on the NovaSeq 6000 platform targeting 50M reads per sample. Raw sequencing data (FASTQ files) obtained from the NovaSeq 6000 was subjected to quality control analysis, including read quality assessment. Real Time Analysis Software (RTA) and NovaSeq Control Software (NCS) (1.8.0; Illumina) were used for base calling. MultiQC (v1.7; Illumina) was used for data quality assessments. Paired-end RNA sequencing reads (150bp long) were trimmed and filtered for quality using Trimgalore v0.6.7 (https://doi.org/10.5281/zenodo.7598955). Trimmed reads were aligned and counted using Spliced Transcripts Alignment to a Reference (STAR) v2.7.9a with the-quantMode GeneCounts parameter against the mm39 mouse genome and GENCODE comprehensive gene annotations (Relsease M31)^40^. ∼50-100 million uniquely mapped reads were acquired per sample. Sample read counts were normalized and differential expression was performed using DESeq2 v1.34.0^39^. Genomic features counted fewer than five times across at least three samples were removed. False discovery rate adjusted for multiple hypothesis testing with Benjamini-Hochberg (BH) procedure p value < 0.05 and log2 fold change >1 was used to define differentially expressed genes. Gene set enrichment analysis (GSEA) was performed using the R package Clusterprofiler^41^ with gene sets from the MSigDB database^42^.

### Organoid bulk RNA-seq analysis

<u>Differential gene expression analysis.</u> RNA-seq raw count data was used as input to the DESeq2(1.44.0) R package DESeq() function. Differential gene expression results were computed, comparing the control and doxycycline treated groups. Variance stabilizing transformation was performed using the DESeq2 varianceStabilizingTransformation() function on the RNA-seq raw count data, and the resulting transformed data (vsd data) was used for down-stream analysis. <u>Differential gene analysis</u> <u>visualization.</u> Differential gene expression results output from DESeq2 were visualized via volcano plots. Matplotlib(3.7.5)’s axes.Axes.scatter() function was used to plot-log10 transformed adjusted p-values over log2 fold change values for each gene. Genes with absolute log2 fold change values >= 2 and adjusted p-value <= 0.05 were highlighted. <u>Heatmap visualization.</u> Scanpy(1.9.3) function scanpy.pl.heatmap() was used to generate the heatmaps. Vsd count matrices output from DESeq2 were used as input to the heatmap function. Heatmap colors represented the gene expression values that were normalized by library size, log1p transformed and z-score scaled. Genes were ordered by ascending differential gene analysis test-statistics on the heatmap. <u>Principal component analysis (PCA) and visualization.</u> For *KrasGNAS* and *GNAS* organoid gene expression, the DESeq vsd count matrices were used to compute the principal components (PC) of the data. For cell line glycan abundance analysis, the mass spectrometry glycosylation profiling data were used. Data matrices were normalized by library size, arcsinh-transformed and z-score scaled for the PCA analysis. scanpy.tl.pca() function was used to perform the PCA, with transformed matrices and other default arguments of the function as inputs. Scatter plots of the first two PC values were generated using the scanpy.pl.pca() function, with data points colored by conditions of interest.

### pySCENIC and gene regulatory network (GRN) inference

To infer the GRN in pyloric metaplasia of the pancreas, we performed SCENIC using pySCENIC functions on a scRNA-seq dataset of murine pancreatitis^20, 43^. This protocol allows the reconstruction of regulons (TF and known target genes) from gene co-expression data, assesses regulon activity in single cells, and can be used to find regulon-enriched cellular clusters. Specifically, we ran v0.11.0 of pySCENIC in a Singularity container built from the Docker Hub image, on ACCRE, Vanderbilt’s High Performance Computing cluster. Following the quality control and feature selection performed in Seurat, we exported its raw counts to a matrix that was then converted to a LOOM file. Alongside a list of 1170 mouse TFs, this gene expression matrix served as input for calculating gene co-expression modules via GRNBoost2. To account for the stochastic nature of GRNBoost2, we calculated the co-expression modules 100 times and then retained only TF/target gene associations that exist in at least 80% of the runs. We then merged the results of these 100 runs as a left outer join operation and averaged the IM values reported for each association. This consensus GRN was then used as input for module pruning, where we filtered out indirect gene targets lacking the cis-regulatory motif associated with the TF. This step used SCENIC’s RcisTarget and ranking databases for motifs (mm9-tss-centered-5kb-7species.mc9nr.feather) in the promoter of the genes [up to 500 base pairs (bp) around the transcriptional start site (TSS)]. The resulting co-expressed TF-target genes are then grouped into regulons. Lastly, the activity of the regulons was computed using SCENIC’s AUCell function, which uses the “area under the curve” (AUC) to calculate whether a subset of the input gene set is enriched within the expressed genes for each cell. These activity data were further binarized (assigned an ON or OFF value, per regulon, per cell) by threshold on the AUC values of the given regulon. Both the AUCell and binarized regulon activity matrices were integrated into Seurat object via the “CreateAssayObject” function, for downstream analysis and visualization^44^. UMAP visuals of the binary and AUC metrics were created from Seurat’s DimPlot() function. Heatmap visuals of binary regulon matrix were performed by ComplexHeatmap R package^45^. Network plots were created from Cytoscape from top 10% based on IM reported for TF/target association from the coexpression modules^46^.

### Statistical Analysis

Statistical analyses and data processing were performed in Image J or Prism (GraphPad, San Diego, CA). Statistical significance was calculated by either 2-tailed unpaired t-tests assuming equal variance or 1-way analysis of variance. Data are expressed as mean ± standard deviation. Figures were generated with Adobe Photoshop and Illustrator.

## RESULTS

### Human IPMN recapitulate pyloric metaplasia

Metaplastic tissues bearing markers of the gastric pylorus have been reported in injury and tumorigenesis in several gastrointestinal organs^20–24^. To evaluate expression of pyloric metaplasia markers in IPMN, we evaluated expression of *MUC5AC* (foveolar pit lineage marker), *TFF2*, *AQP5*, and *CD44* (SPEM markers), which produces splicing variant CD44v9, in published RNA-seq datasets derived from patient samples. Previously, Bernard et al., generated a single cell RNA-sequencing (scRNA-seq) dataset composed of epithelium and stroma from either low-grade or high-grade IPMN (n = 2 each) or IPMN associated PDAC (n = 2)(**Figure S1A-C**)^35^. Analysis of this dataset for molecular markers of IPMN subtype revealed widespread expression of *MUC5AC* throughout the epithelium of all samples, with *MUC6* and *MUC2* differentiating gastric-from intestinal-type IPMN, respectively (**Figure S1D**). Expression of *TFF2* and *AQP5* was enriched in the epithelium whereas *CD44* was identified in both the epithelium and stroma (**Figure S1E**). To evaluate a second dataset, we interrogated bulk RNA-seq generated by Maurer et al., of laser capture dissected epithelium or stroma from patient IPMN (n = 19), PanIN (n = 26), or PDAC (n = 197 epithelium, 124 stroma)^37, 38^. Like our analysis of IPMN scRNA-seq data, we identified expression of *MUC5AC* and *AQP5*, enriched in the epithelium of all disease states, and *CD44,* in both epithelial and stromal populations (**Figure 1A**). Finally, we examined expression of a SPEM gene signature generated from our previously published murine pancreatitis scRNA-seq dataset, in the Maurer dataset. We identified enrichment of several additional markers in pre-invasive PanIN and IPMN, as compared to PDAC (**Figure S2**)^20^.

**Figure 1.**
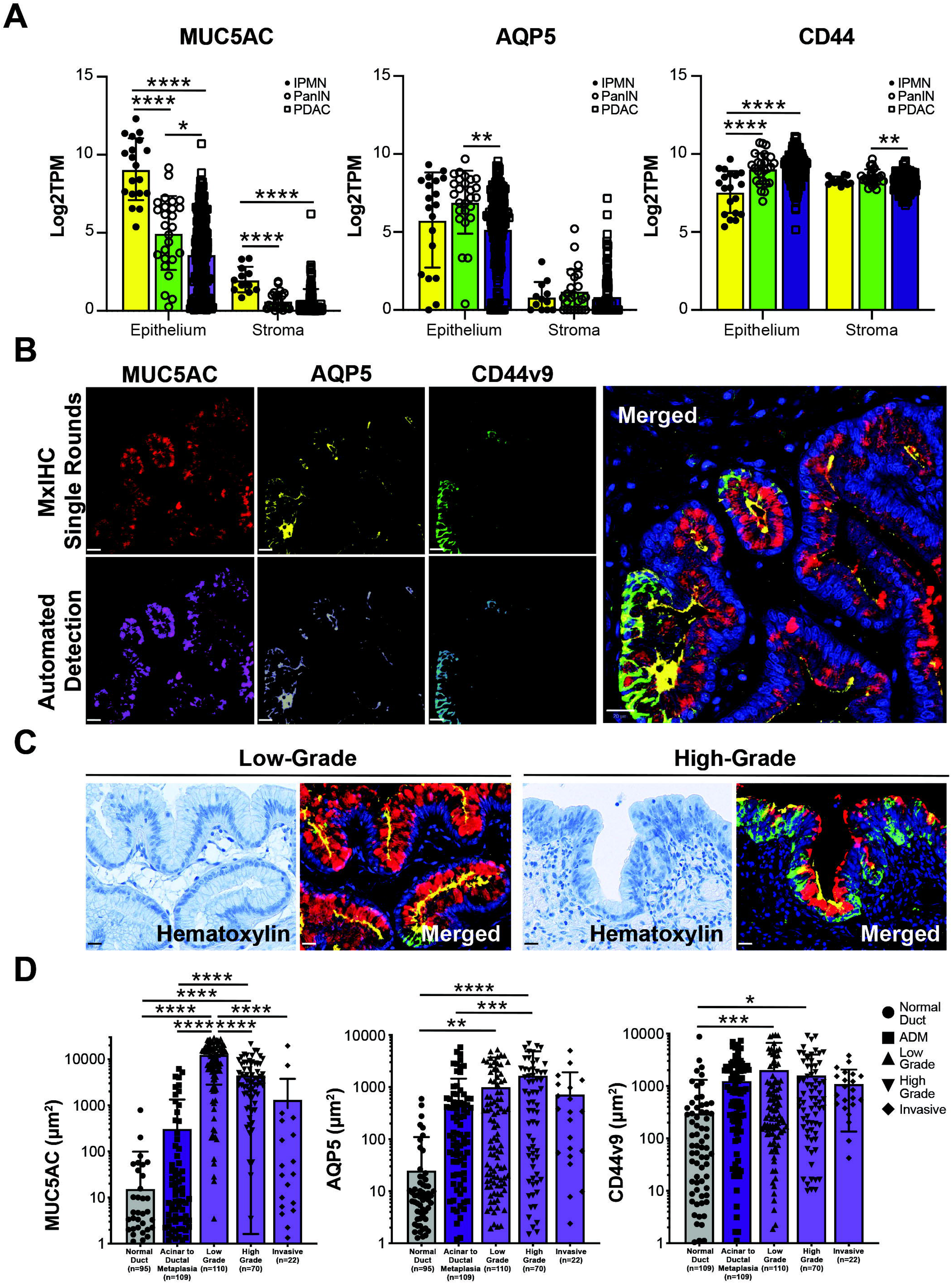
Human IPMN are characterized by pyloric metaplasia. (**A**) Bar plots comparing expression (Log2TPM) of microdissected epithelium and stroma from Maurer et al.^37^ from IPMN (n = 19), PanIN (n = 26), and PDAC (n = 197 epithelium, 124 stroma) for *MUC5AC*, *AQP5*, or *CD44*. (**B**) Pseudo-colored immunohistochemical staining for pyloric metaplasia markers MUC5AC (red), AQP5 (yellow) or CD44v9 (green), top row, and automated detection of signal (bottom row) by QuPath to merge MxIHC data^34^. (**C**) Examples of hematoxylin staining or merged MxIHC staining of low grade (left) or high grade (right) IPMN. Scale bars, 20 mm. (**D**) Quantification of staining in (B-C) for 41 IPMN patients including normal ducts (n = 95-109), acinar to ductal metaplasia (n = 109), low grade IPMN (n = 110), high grade IPMN (n = 70), and invasive IPMN (n = 22). *, p < 0.05; **, p < 0.01; ***, p < 0.005; ****, p < 0.001.

To confirm expression of pyloric metaplasia markers in patient IPMN samples, we conducted multiplex immunohistochemistry (MxIHC) on a cohort of 41 patients (**File S1**). Serial staining was conducted for MUC5AC, AQP5, and CD44v9 and expression scored in multiple tissue compartments within each sample. Signal for each stain was automatically detected using QuPath and overlaid to generate merged images (**Figure 1B**)^34^. Altogether, expression was evaluated in normal ducts (n = 95-109 regions of interest, ROIs), acinar to ductal metaplasia (ADM, n = 109 ROIs), low-grade IPMN (n = 110 ROIs), high-grade IPMN (n = 70 ROIs), and foci of invasive PDAC (n = 22 ROIs). All markers were detected; interestingly, protein localization changed between low-and high-grade IPMN (**Figure 1C**). Consistent with previous reports; we identified a significant increase in MUC5AC expression between normal ducts and IPMN and a significant decrease in expression with progression from low to high-grade, invasive IPMN^35^. Expression of both AQP5 and CD44v9 increased with disease progression, reaching significance in low and high-grade IPMN (**Figure 1D, S3A-B**). Stratifying IPMN by molecular subtype (gastric, intestinal, pancreatobiliary-like), we identified pyloric metaplasia in all subtypes with no significant difference in expression of MUC5AC or AQP5 between gastric and intestinal type IPMN (**Figure S3C**). Collectively, these data demonstrate that a significant increase in pyloric metaplasia marker expression accompanies IPMN formation in patients.

### Mutant GNAS drives a pyloric phenotype

While PanIN and IPMN have both been shown to express oncogenic *KRAS*, most IPMN also express mutant *GNAS*^11^. Previous studies have shown that *GNAS^R201C/H^* mutations inhibit PDAC aggressiveness^13, 14^. To determine if *GNAS^R201C^* drives an indolent phenotype by amplifying a pyloric phenotype, we used two cell lines (4838 and C241) harboring a *Kras^G12D^* mutation that conditionally express human *GNAS^R201C^*upon doxycycline (DOX) treatment (**Figure 2A**)^13^. Consistent with previous studies, we identified a significant decrease in cancer cell invasion in both cell lines with DOX (**Figure 2B**). Correspondingly, we saw an increase in expression of epithelial markers *Epcam* and *Cldn2* and a decrease in mesenchymal markers *Fn1* and *Zeb1* (**Figure S4**) by qRT-PCR, suggesting a mesenchymal to epithelial transition (MET). To identify potential mechanisms by which *GNAS^R201C^* inhibits tumor cell aggressiveness, the 4838 and C241 cell lines were treated +/-DOX and underwent RNA sequencing (n = 6, 3 biological replicates/condition) (**Figure 2C-E, Figure S4, File S2-3**). As expected, there was a significant increase in human (hu*GNAS*), but not murine *Gnas* (mu*Gnas*) expression (**Figure 2D**). Significant differences were identified between the two cell lines, likely due to biological heterogeneity or differences in tumor suppressor mutations (**Figure 2E, S4**); however, both cell lines showed an increase in expression of pyloric markers (*Gkn1*, *Aqp5*, *Tff2*) and mucin genes (*Muc5ac*) with induction of *GNAS^R201C^*expression (**Figure 2F)**. Consistent with our assessment, gene set enrichment analysis (GSEA) identified a decrease in expression of genes associated with EMT and EGFR signaling, as well as an increase in expression of genes associated with early gastric cancer (**Figure S4F-G**).

**Figure 2.**
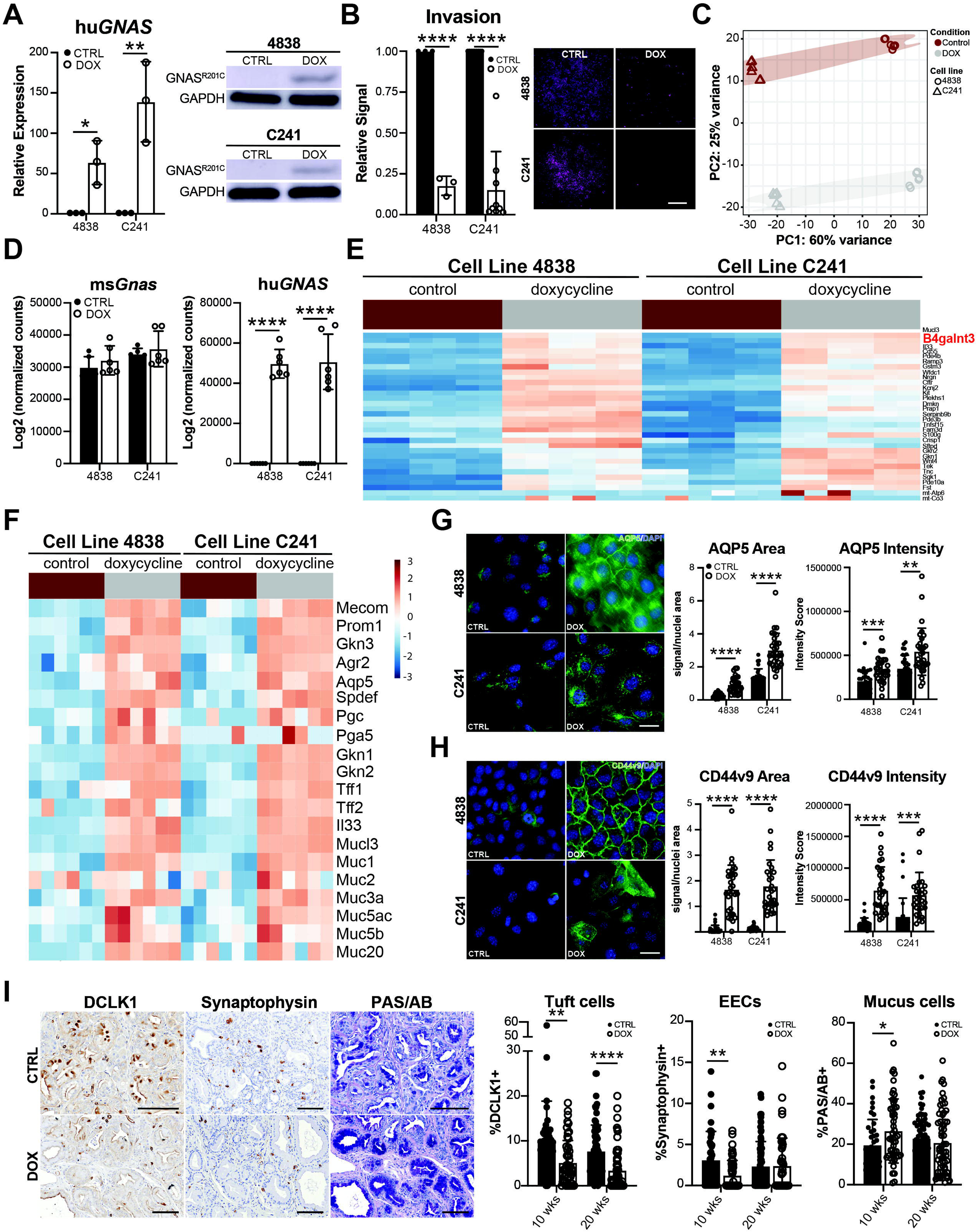
**Mutant GNAS drives a pyloric metaplasia program**. (**A**) qRT-PCR (n = 3) or western blot for human *GNAS* (hu*GNAS*) in 4838 and C241 cell lines treated +/- doxycycline (DOX). (**B**) Representative images and quantification of invasion for both cell lines treated +/- DOX. Scale bar, 1 mm. (**C**) Principal component analysis (PCA) of RNA-seq conducted on both cell lines +/- DOX (n = 6, 3 biological replicates). (**D**) Bar plots of murine (ms) or human (hu) *GNAS* expression, determined by RNA-seq, in both cell lines. (**E**) Heatmap of top upregulated genes in both cell lines with DOX. (**F**) Heatmap of pyloric metaplasia marker gene expression in both cell lines +/- DOX. (**G**) Immunofluorescence and quantification of signal area and intensity for AQP5 and (**H**) CD44v9 in both cell lines +/- DOX (n = 3 biological replicates). Scale bars, 25 mm. (**I**) IHC and quantification of tuft marker DCLK1, EEC marker synaptophysin, or mucin staining by PAS/AB in *KrasGNAS* pancreata from mice fed a control or DOX diet. Scale bars, 100 mm. *, p < 0.05; **, p < 0.01; ***, p < 0.005; ****, p < 0.001.

To confirm gene expression changes for select markers at the protein level, we conducted immunofluorescence (IF) on 4838 and C241 cells +/- DOX. As shown in **Figure 2G-H**, we saw a significant increase in both AQP5 and CD44v9 in overall expression and intensity. To determine whether *GNAS^R201C^* preferentially drives mucus cell production *in vivo*, we next evaluated cell type abundance in *KrasGNAS* mice. Pancreata were collected from *KrasGNAS* mice treated with or without DOX for either 10 or 20 weeks (n = 3 mice/condition)^13^. H&E were then evaluated by a pathologist (VQT) for disease progression and cyst formation. When combined, the two groups were found to be equivalent in terms of lesion grade, however, DOX treatment was found to enhance cyst formation (**Figure S5A**), as previously described^13^. Regions of cyst/IPMN and PanIN lesions were then identified by H&E (∼3 regions of each/slide), with most mice harboring both (up to 6 areas identified/mouse). Previously, we showed that metaplasia in the pancreas results in the *de novo* formation of several differentiated cell types, such as tuft, enteroendocrine (EEC), and mucus-producing cells^20, 47–49^. Serial sections were stained for tuft cell marker DCLK1^47^, EEC marker synaptophysin^50, 51^, and mucins (PAS/AB) and quantified. As shown in **Figure 2I**, *GNAS^R201C^* expression was sufficient to significantly decrease tuft (9.9%, 45 lesions vs. 5.2%, 52 lesions) and enteroendocrine cell numbers (3.1%, 42 lesions vs. 1.2%, 52 lesions) while significantly increasing mucin production (19.6%, 40 lesions vs. 26.6%, 52 lesions) in low grade lesions, consistent with our analysis of human data (**Figure S5B, File S4**). Collectively, these data demonstrate that *GNAS^R201C^*shifts cells away from aggressive behavior and towards a mucinous, pyloric phenotype, potentially limiting tumor cell aggressiveness by driving a program of differentiation.

### Identification of master regulators of pyloric metaplasia

Previously, we showed through scRNA-seq, immunostaining, and electron microscopy that pancreatitis is characterized by a cell population resembling SPEM^20^. To identify master regulator transcription factors driving this phenotype, we performed a PyScenic-based Regulon analysis on our pancreatitis scRNA-seq dataset (∼13,000 cells), which identifies candidate factors by expression of known downstream target genes (**Figure 3A**). Among the candidate regulators, we identified the transcription factor *Spdef*, which has been shown to be the master regulator of mucin-producing goblet cell formation in the stomach, intestines, and lung (**Figure 3B**)^52–54^. Consistent with a role for driving mucin production, we found that *GNAS* mutations in human cell lines are associated with both elevated mucin and *SPDEF* expression (**Figure 3C, S5C**) and that *GNAS^R201C^* expression is sufficient to drive *Spdef* expression in *Kras;GNAS* cell lines (4838 and C241) (**Figure 3E**). Among the top 10% of predicted *Spdef* target genes, we identified several SPEM markers (*Gkn3*, *Muc6*) as well as additional transcription factors predicted to drive this phenotype (*Creb3l1*, *Creb3l4*) (**Figure 3D**, **File S5**). Consistent with *Spdef* being induced by mutant GNAS, *GNAS^R201C^* was sufficient to increase expression of both *Creb3l1* (4838 cells) and *Creb3l4* (4838 and C241 cells) (**Figure 5E**). Regulon analysis predicted that both *Creb3l1* and *Creb3l4* regulate expression of canonical SPEM markers (*Aqp5*, *Tff2*, *Muc6*, among others) as well the expression of each other (*Spdef*, *Creb3l1* and *Creb3l4*) (**Figure S6-7, File S6, File S7**). To compare these data to *GNAS^R201C^*-induced changes in gene expression, we first overlaid the DOX-on gene signatures from both the 4838 and C241 cell lines onto the pancreatitis scRNA-seq dataset and identified enrichment in the SPEM cluster (**Figure 3F-G**); expression patterns closely resembled that for *Spdef* (**Figure 3B**). We next examined changes in expression of *Spdef*, *Creb3l1*, and *Creb3l4* target genes with DOX treatment and *GNAS^R201C^* expression. We identified an increase in target genes in both cell lines for all three transcription factors, including SPEM markers (**Figures S8**, **S9**), suggesting that *GNAS^R201C^* drives expression of a pyloric metaplasia program through the activity of one or more of these master regulator transcription factors.

**Figure 3.**
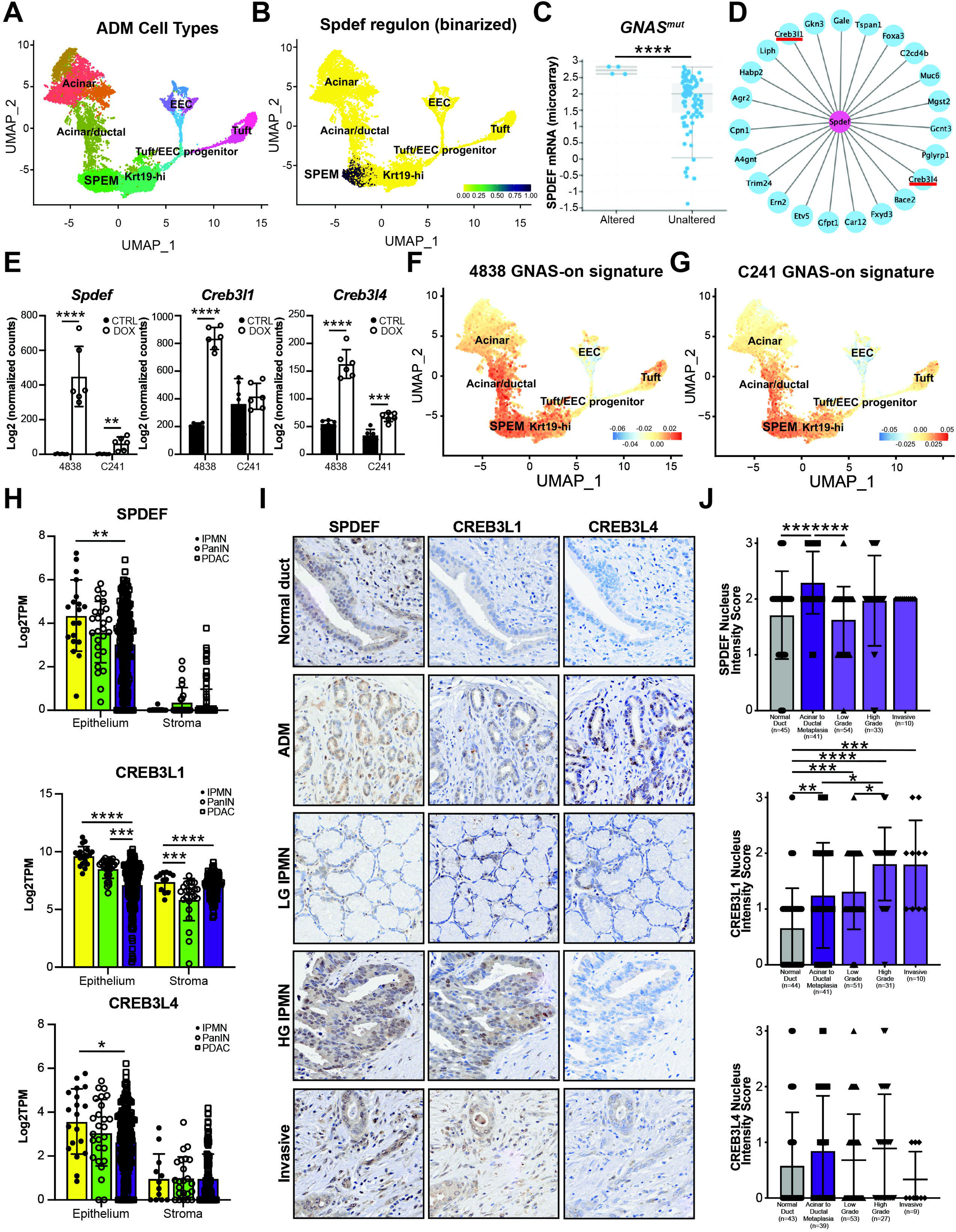
**Identification of master regulator transcription factors of pyloric metaplasia**. (**A**) UMAP of scRNA-seq data from a murine model of pancreatitis showing the formation of a gastric SPEM^20^. (**B**) Binarized activity of *Spdef* predicted by regulon analysis overlayed on the UMAP from (A). (**C**) *SPDEF* expression in PDAC tumors from cBioPortal with wild type (n = 96) or mutant GNAS^R201C/H^ (n = 4). (**D**) Plot of *Spdef* targets predicted by regulon analysis identifying gastric SPEM markers (*Gkn3*, *Muc6*) and transcription factors *Creb3l1* and *Creb3l4*. (**E**) Barplots showing increased expression of *Spdef*, *Creb3l1*, and *Creb3l4* in IPMN cell lines with DOX treatment and GNAS^R201C^ expression. (**F**) DOX/GNAS-on gene signatures from IPMN cell line 4838 or (**G**) C241 overlayed on the UMAP in (A) showing overlap with predicted *Spdef* activity. (**H**) Barplots comparing expression (Log2TPM) of microdissected epithelium and stroma from Maurer et al.^37^ from IPMN, PanIN, and PDAC for *SPDEF*, *CREB3L1*, and *CREB3L4*. (**I-J**) Representative IHC and quantification for SPDEF, CREB3L1, and CREB3L4 conducted on 23 IPMN patient specimens including normal ducts (n = 95-109), acinar to ductal metaplasia (n = 109), low grade IPMN (n = 110), high grade IPMN (n = 70), and pancreatic cancer (Invasive, n = 22). *, p < 0.05; **, p < 0.01; ***, p < 0.005; ****, p < 0.001.

To investigate the relevance of SPDEF, CREB3L1, and CREB3L4 to human IPMN and PDAC, we next examined expression in patient samples. First, we interrogated the bulk RNA-seq dataset generated by Maurer et al., of IPMN (n = 19), PanIN (n = 26), and PDAC (n = 197 epithelium, 124 stroma)^37, 38^. Expression of all three transcription factors was elevated in the epithelium of all disease states, as well as elevated *CREB3L1* in the stroma (**Figure 3H**). To examine protein expression, we next conducted IHC on 23 tissue samples collected from patients with IPMN +/-associated PDAC and scored expression. Regions of interest encompassed normal ducts (n = 43-45), acinar to ductal metaplasia (ADM, n = 39-41), LG IPMN (n = 51-54), HG IPMN (n = 27-33), and invasive PDAC (n = 9-10). Low expression of all three transcription factors was detected in a portion of normal ducts with expression increasing in ADM and reaching significance for SPDEF and CREB3L1 (**Figure 3I-J**). While CREB3L4 expression remained low throughout IPMN to PDAC progression, SPDEF remained high, though expression did not significantly increase between LG IPMN and PDAC. CREB3L1 expression, though, did significantly increase with disease progression from normal ducts through IPMN to PDAC (**Figure 3I-J**). Collectively, these data suggest a possible role for SPDEF and CREB3L1 as master regulators of a pyloric metaplastic phenotype in GNAS mutated IPMN.

### Mutant GNAS drives major shifts in glycan species abundance

Consistent with GNAS^R201C^ driving a pyloric metaplasia phenotype strongly defined by changes in mucin production, we noted significant upregulation of N-acetyl*-*galactosaminyltransferase *B4galnt3*, which synthesizes LacdiNAc glycans (**Figure 2E**) at the expense of Lewis epitopes^55^. Glycans are carbohydrate moieties that modify cell surface receptor function and are a predominant feature of mucins^56^. Specific glycans (e.g. acidic Lewis epitopes like CA19-9 and 3’-Sulfo-Le^A/C^) are used clinically as biomarkers to detect high-grade dysplasia and cancer in IPMNs and to monitor therapeutic response and recurrence in PDAC^57,58^. Forced expression of these Lewis epitopes in murine models suggest that such glycans are pathogenic and promote the development of pancreatic adenocarcinoma^57^. This is contrast to LacdiNAc glycans which have not been associated with high-grade dysplasia and cancer.

To examine global changes in glycosyltransferase expression, we interrogated our dataset for expression of 137 different enzymes (**Figure S10A**), and identified significant changes in several, including upregulation of *Mgat4a*, which catalyzes the transfer of GlcNAc to generate branched glycan structures, and *Galnt2*, which initiates mucin-type O glycosylation by transferring a GalNac to serine and threonine sidechains, consistent with an increase in mucin gene expression (**Figure 4A-B**). To determine if these changes are directly due to *GNAS^R201C^* or a consequence of differentiation, we generated organoids from *KrasGNAS* pancreata lacking disease. Several lines (n = 3 mice) were propagated in culture, treated +/- DOX to induce *GNAS^R201C^*, and underwent RNA-seq (**Figure 4C, File S8**). Consistent with *GNAS^R201C^* driving a pyloric phenotype, we identified significant upregulation of *Muc5ac* (**Figure 4D**), in addition to glycosyltransferases *B4galnt3*, *Galnt2*, and *Mgat4a* (**Figure 4E-F**) in DOX-treated organoids in contrast to non-DOX-treated organoids. Similar gene expression changes were identified in *GNAS* organoids +/- DOX lacking expression of *Kras^G12D^* (**Figure S10B-E**). Collectively, these data show that *GNAS^R201C^* drives a pyloric phenotype (in precancer and PDAC), which includes significant changes in glycosyltransferase expression.

**Figure 4.**
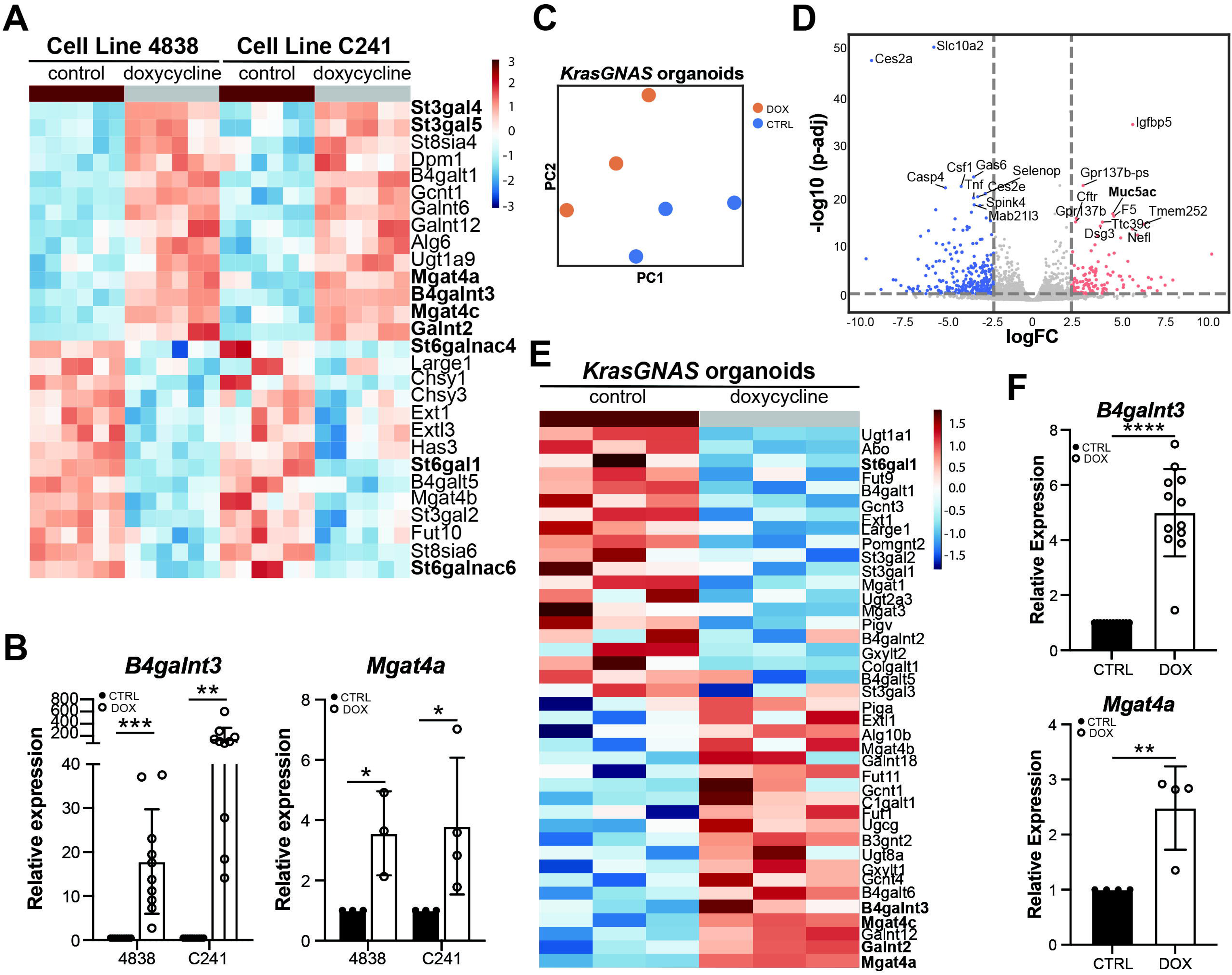
Mutant GNAS drives significant changes in glycosyltransferase gene expression. (**A**) Heatmap of select glycosyltransferase gene expression in 4838 and C241 cells treated +/- DOX. (**B**) qRT- PCR for either *B4galnt3* (n = 10) or *Mgat4a* (n = 3-4) in both cell lines +/- DOX. (**C**) PCA of RNA-seq conducted on organoids generated from *KrasGNAS* mice treated +/- DOX. (**D**) Volcano plot of differentially expressed genes in *KrasGNAS* organoids +/- DOX, highlighting pyloric metaplasia marker *Muc5ac*. (**E**) Heatmap of select glycosyltransferase gene expression in *KrasGNAS* organoids treated +/- DOX. (**F**) qRT- PCR for either *B4galnt3* (n = 4 biological, 3 technical replicates) or *Mgat4a* (n = 4 biological replicates) in both cell lines +/- DOX. *, p < 0.05; **, p < 0.01; ***, p < 0.005; ****, p < 0.001.

To determine if these changes in glycosyltransferase gene expression translate to changes in glycan abundance with *GNAS^R201C^*, cell lines 4838 and C241 were treated +/- DOX then stained with lectins recognizing both LacdiNAcs (*B4galnt3*) and GlcNAcs (*Mgat4a* and/or *Gcnt2*, which bias towards GlcNAc high branching glycans). As shown in **Figure 5A-B**, we saw a significant increase in both area and intensity of *Wisteria floribunda* (WFA) staining in both cell lines with DOX treatment, reflecting an increase in LacdiNAc glycans. Similarly, we saw an increase in GlcNAc modifications by lectin *Griffonia simplicifolia* (GSII) in both cell lines with *GNAS^R201C^* expression. Interestingly, we saw a significant decrease in intensity of Das-1 staining with DOX, which recognizes 3’ sulfated Lewis A/C glycans (3’-sulfo-Le^A/C^), a marker for high grade dysplasia and PDAC, consistent with *GNAS^R201C^* driving a less aggressive phenotype (**Figure 5C**)^58, 59^. To confirm these changes and explore global changes in glycosylation induced by *GNAS^R201C^*, we performed glycosylation profiling by mass spectrometry (**Figure 5D-E**, **File S9**). While minor changes in O glycan species were identified with DOX treatment (**Figure S11**), we saw a significant shift in N glycan abundance in both cell lines (**Figure 5F-G**, **S12**). Among these changes, we identified a significant increase in LacdiNAc species, consistent with WFA lectin staining (**Figure 5H**). We also identified a loss of Lewis^A/X^ abundance in both cell lines (N5H4F1), consistent with a loss of Das-1 staining and a less aggressive phenotype (**Figure 5I**). To confirm these changes with *GNAS^R201C^* expression *in vivo*, we performed imaging glycan mass spectrometry on pancreas tissue sections from *KrasGNAS* mice fed +/- DOX chow for 22 weeks. While less sensitive than our cell line analysis, we were able to identify a significant increase in glycan abundance in PanIn and IPMN, as compared to acinar tissue, in fucosylation, sialyation, and predicted LacdiNAcs (**Figure S13**). Altogether, these data demonstrate that *GNAS^R201C^* drives synthesis of LacdiNAcs which is associated with a loss of Lewis epitopes^55^. These changes in glycan species/abundance may represent a mechanism by which *GNAS^R201C^* inhibits tumor aggressiveness.

**Figure 5.**
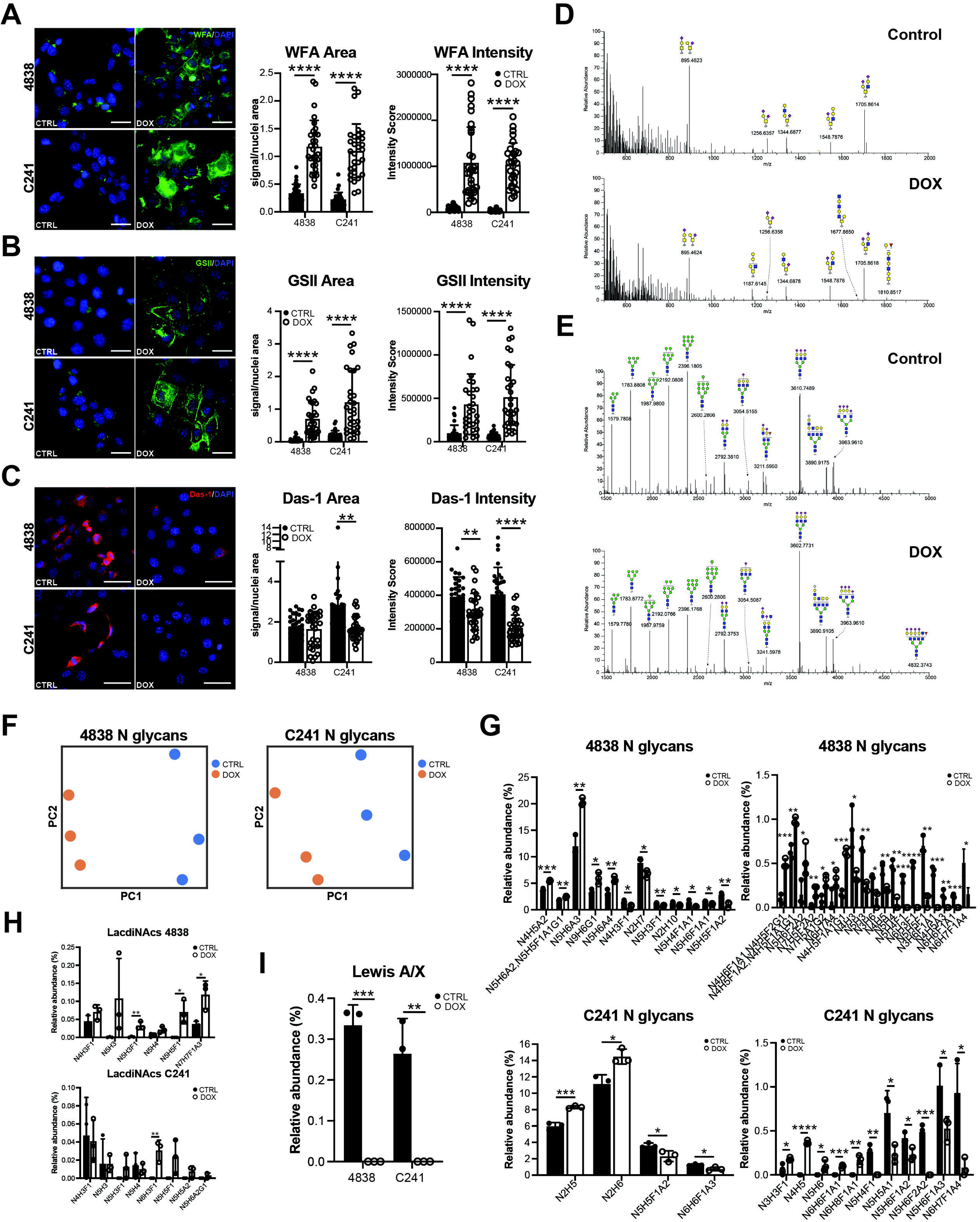
Mutant GNAS drives significant changes in N glycosylation. IF and quantification of (**A**) lectin WFA recognizing LacdiNAcs (**B**) lectin GSII recognizing GlcNAcs or (**C**) Das-1 antibody staining recognizing 3’-sulfo-Le^A/C^ in 4838 and C241 cells +/- DOX. Scale bars, 50 mm. Spectra of (**D**) O- or (**E**) N- glycans identified by mass spectrometry in 4838 cells treated +/- DOX. (**F**) PCA of N glycan changes in 4838 or C241 cells +/- DOX. (**G**) Bar plots highlighting significant changes in N glycans, (**H**) LacdiNAcs, or (**I**) Lewis^A/X^ antigen in both cell lines +/- DOX. *, p < 0.05; **, p < 0.01; ***, p < 0.005; ****, p < 0.001.

### GNAS^R201C^-induced glycan changes drive an indolent phenotype

Our regulon analysis identified SPDEF and CREB3L1 as possible transcriptional regulators of the *GNAS^R201C^*- induced pyloric phenotype. To determine if either transcription factor is responsible for *B4galnt3*-driven LacdiNac deposition in response to *GNAS^R201C^*, we performed knockdown experiments using small interfering RNA (siRNA). In response to treatment with siRNA targeting *Spdef*, we identified a significant decrease in expression of *Spdef*, pyloric marker *Gkn1*, and *B4galnt3* in both cell cells by qRT-PCR (**Figure 6A**). We also saw a reduction in *GNAS* expression and in epithelial markers *Epcam* and *Cldn2*, without a corresponding increase in mesenchymal marker *Zeb1* (**Figure S14A-B**). Consistent with a loss of *B4galnt3*, we saw a significant decrease in WFA lectin staining in area and intensity in both cell lines with *Spdef* siRNA as compared to control (**Figure 6B, S14D-E**). In response to treatment with siRNA targeting *Creb3l1*, we identified a significant decrease in expression of *Creb3l1*, as well as pyloric marker *Gkn1*, and *B4galnt3* in both cell cells by qRT-PCR (**Figure 6C**). Similar to results obtained with *Spdef* siRNA, we again saw a decrease in *GNAS*, *Epcam*, and *Cldn2* (but no increase in *Zeb1*) as well as a loss of WFA area and intensity with *Creb3l1* siRNA as compared to control in both cell lines (**Figure 6C-D, S14F-H)**. These data suggest that transcriptional regulators of the *GNAS^R201C^*pyloric phenotype also regulate LacdiNAc abundance.

**Figure 6.**
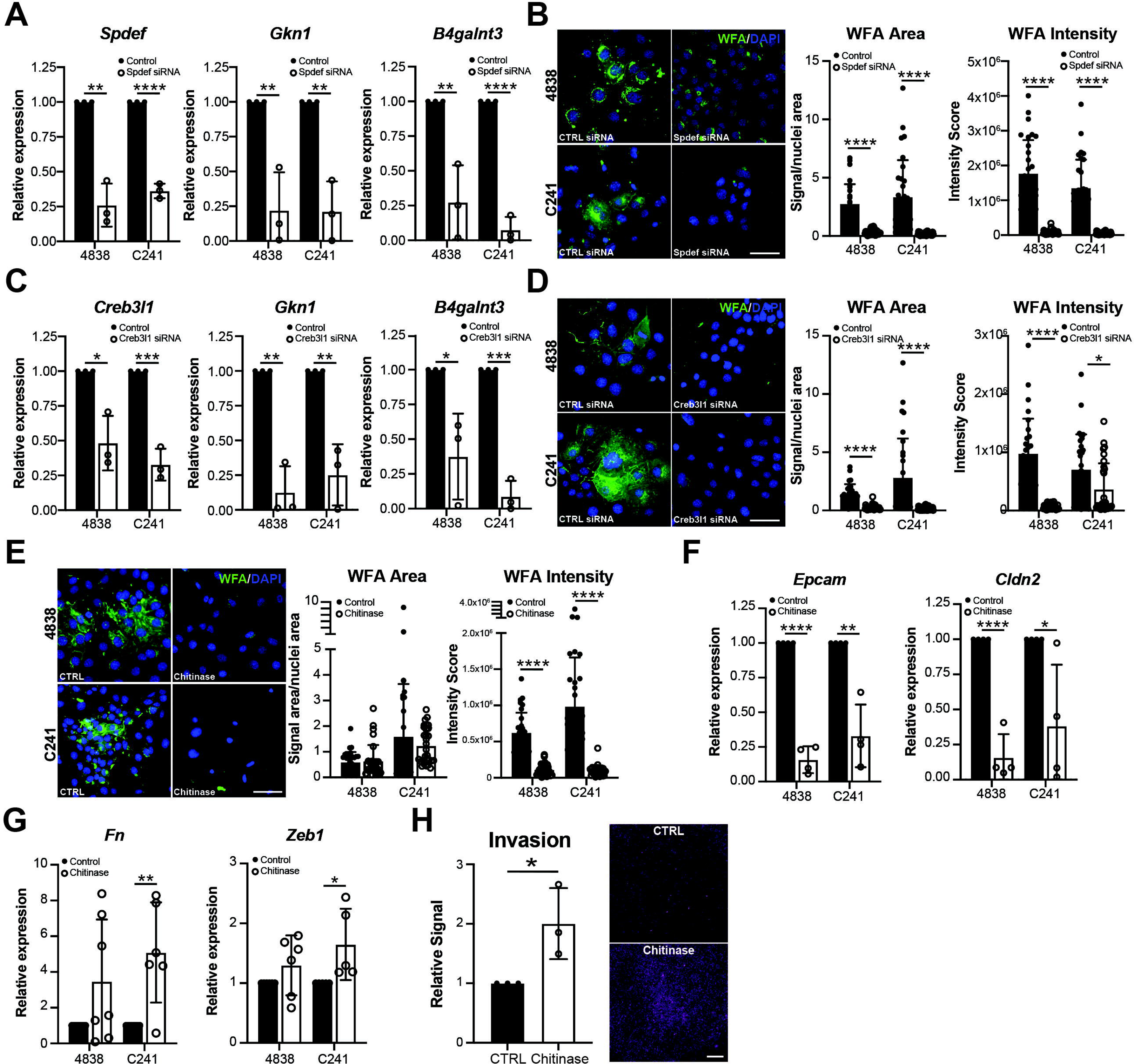
Mutant GNAS drives an indolent phenotype through LacdiNAc deposition. (**A**) Bar plots of *Spdef*, *Gkn1*, or *B4galnt3* expression in 4838 or C241 cells treated with DOX and either control or *Spdef* siRNA, determined by qRT-PCR. (**B**) IF and quantification of lectin WFA in both cells lines treated with DOX and either control or *Spdef* siRNA. Scale bar, 50 mm. (**C**) Bar plots of *Spdef*, *Gkn1*, or *B4galnt3* expression in 4838 or C241 cells treated with DOX and either control or *Creb3l1* siRNA, determined by qRT-PCR. (**D**) IF and quantification of lectin WFA in both cells lines treated with DOX and either control or *Creb3l1* siRNA. Scale bar, 50 mm. (**E**) IF and quantification of lectin WFA in both cells lines treated with DOX and either control or chitinase. Scale bar, 50 mm. (**F**) Bar plots of *Epcam* or *Cldn2* expression in 4838 or C241 cells treated with DOX and either control or chitinase, determined by qRT-PCR. (**F**) Bar plots of *Fn* or *Zeb1* expression in 4838 or C241 cells treated with DOX and either control or chitinase, determined by qRT-PCR. (**H**) Quantification and representative images of invasion for C241 cells treated with DOX and either control or chitinase. Scale bar 1 mm. *, p < 0.05; **, p < 0.01; ***, p < 0.005; ****, p < 0.001.

Previous studies have found that chitinase, an enzyme known to break down chitin, a non-mammalian structural protein found in fungi cell walls and the exoskeletons of crustaceans and insects, also hydrolyzes the glycosidic bond in LacdiNAcs (GalNAcβ1-4GlcNAc)^60^. To determine if chitinase and LacdiNAc cleavage impacts the *GNAS^R201C^*-induced phenotype, we treated DOX-incubated 4838 and C241 cells with chitinase. As shown in **Figure 6E**, we identified a significant decrease in WFA lectin intensity, consistent with LacdiNAc cleavage. Interestingly, we saw a significant decrease in epithelial markers *Epcam* and *Cldn2* in both cell lines with and an increase in mesenchymal markers *Zeb1* and *Fn1* in C241 (**Figure 6F-G**). Consistent with a reversal of the *GNAS^R201C^*-induced MET phenotype, we saw an increase in cancer cell invasion with chitinase treatment in C241 cells (**Figure 6H**). Altogether, these data suggest that LacdiNAc deposition represents a mechanism by which *GNAS^R201C^* inhibits cancer cell aggressiveness. Further studies are required to determine what cell surface receptors are modified with LacdiNAcs and how this impacts their activity.

### Glycans may be used to distinguish low and high grade IPMN

In our *in vitro* and *in vivo* analyses, we identified distinct glycosylation changes associated with low grade disease and the *GNAS^R201C^*-driven, indolent phenotype (WFA+GSII+). To correlate these glycosylation findings with IPMN pathogenesis in patients, we performed co-IF for WFA and GSII, as well as the 3-sulfo-Le^A/C^ marker Das-1, which has previously been shown to label high-grade and invasive disease, in 27 patient IPMNs. Consistent with our *in vitro* data, LacdiNAc-bound WFA and GSII were largely expressed in metaplasia and low-grade IPMN (**Figure 7A-B**). As in prior reports, Das-1 was enriched in high-grade and invasive IPMN (**Figure 7C**)^58^. We noted significant expression of glycans within the intraluminal mucin, correlating to tumor grade (**Figure 7A-F**). Furthermore, combined staining of all markers highlighted small niches of morphologically high-grade cells within mostly low-grade IPMN (**Figure 7H-I**). Collectively, human glycan abundance largely recapitulated our experimental findings and may represent a new strategy to risk stratify IPMN patients.

**Figure 7.**
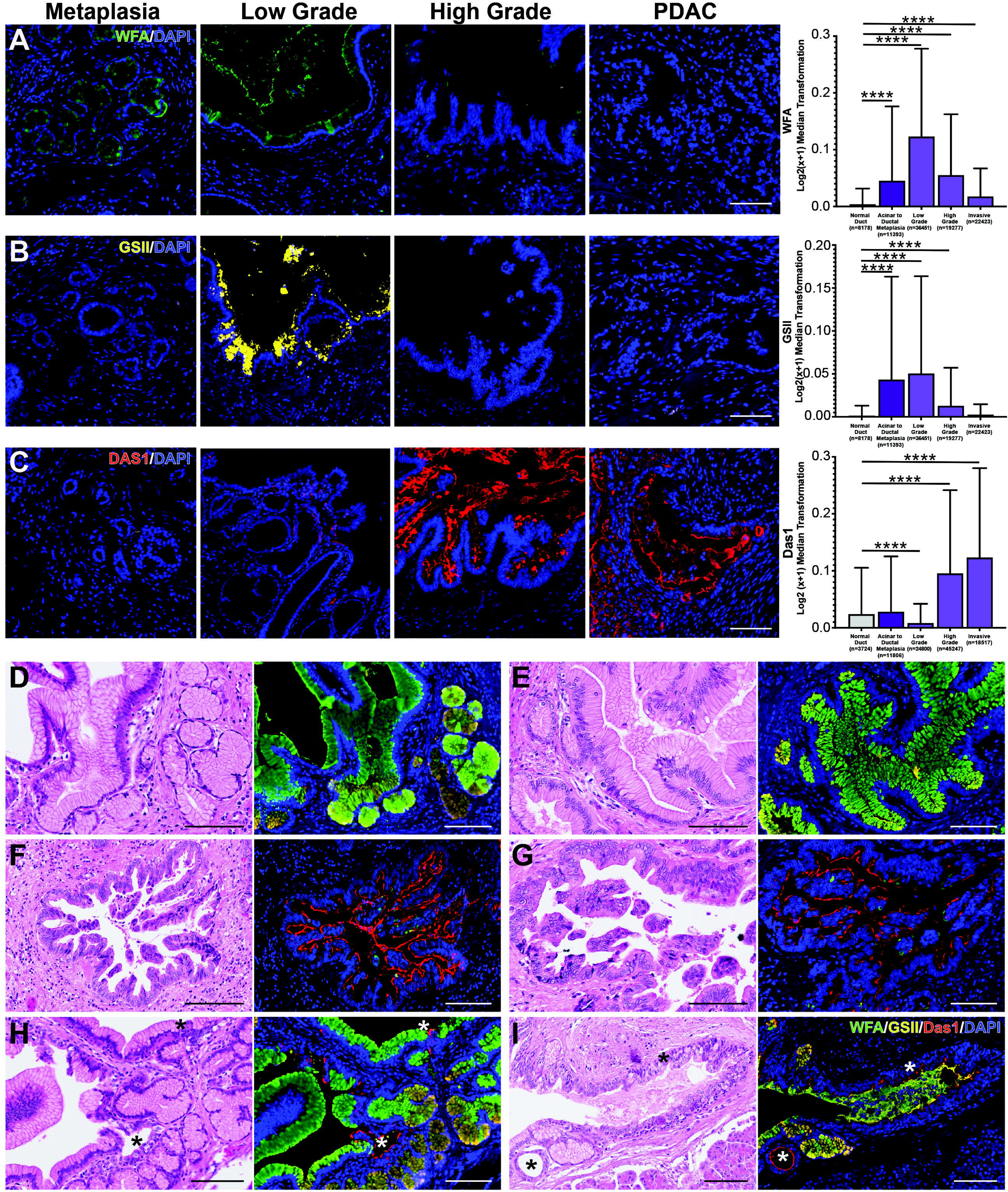
Glycans distinguish low-from high-grade IPMN and PDAC. IF and quantification of (**A**) lectin WFA recognizing LacdiNAcs (green), (**B**) lectin GSII recognizing GlcNAcs (yellow), or (**C**) Das-1 recognizing 3’-sulfo-Le^A/C^ (red) in either metaplasia, low grade or high grade IPMN, or PDAC in 41 IPMN patient samples. Scale bars, 100 mm. Representative images of (**D-E**) low grade IPMN, (**F-G**) high grade IPMN, and (**H-I**) low grade IPMN with foci of high-grade cells co-stained for WFA (green), GSII (yellow), and Das-1 (red). Scale bars, 100 mm. ****, p < 0.001.

## DISCUSSION

PanIN and IPMN, the two most prevalent PDAC precursor lesions, are defined by mucin-producing cells harboring defining genetic alterations. The involvement of pyloric metaplasia has been described previously in murine pancreatitis and PanIN and now in human and murine IPMNs^20, 21^. Pyloric metaplasia-defining MUC5AC, CD44v9, and AQP5 are strongly expressed in acinar-to-ductal metaplasia and their co-expression is identified in dysplastic stages of IPMN progression (**Figure 1**). Furthermore, PanIN and IPMN pyloric metaplasia cells harbor similar whole-transcriptomic signatures, reciprocating gastric SPEM and foveolar pit cell lineages and raising the possibility of a conserved program between these organs^20, 21^. The identification of similar processes of injury and repair between gastrointestinal organs could lead to the discovery of targetable pathways for multiple inflammatory or pre-malignant conditions.

IPMNs differ clinically from PanIN through their capacity to be detected by routine imaging due to their larger size and cystic nature, filled with mucin. Some studies suggest that adenocarcinomas associated with IPMN have significantly better survival trends when compared to PanIN-associated PDAC, even with propensity score matching^61, 62^. Consistent with this and previous studies, we show here that GNAS^R201C^ drives a more indolent phenotype, suggesting that GNAS mutations may explain this disparity^13^. We discovered that both oncogenic KRAS and GNAS drive pyloric metaplasia signatures, yet oncogenic GNAS^R201C^ amplifies a mucinous phenotype, consistent with a role described for GNAS^R201H^ ^20, 63^. A recent clinical study showed that mucinous tumors from multiple tissue types are enriched for GNAS variants, suggesting a conserved role for GNAS mutations in tumorigenesis^64^. How these changes in epithelial heterogeneity affect disease progression remains to be determined.

Through regulon analyses and siRNA-mediated knockdown, we found that mutant GNAS likely drives these changes through a master-regulator SPDEF-CREB3L1 axis. Indeed, these markers increase in expression as early as the acinar-to-ductal metaplasia phase, and CREB3L1 expression continues to increase with disease progression. SPDEF activity is a conserved process identified in the gastric, intestinal, and pulmonary epithelial response to injury and dysplasia, and was recently shown to drive a mucinous phenotype in PanIN^52–54, 65^. This further corroborates that not only is pyloric metaplasia conserved between organs, but also that distinct master regulators underly these processes and the transition between cell states. while some studies have been conducted looking at the functional role for individual pyloric markers in disease progression^31^, it remains to be determined if SPEM and the subsequent addition of a foveolar pit lineage, or pyloric metaplasia, is protective or reflects a detectable sign of disease progression. Future studies assaying for this program may be useful in staging IPMN, while functional studies on the overall role of this program may determine any benefit inducing this differentiation program in advanced disease.

In addition to driving mucus cell formation through SPDEF-CREB3L1, we found that this axis drives major shifts in N-glycan species abundance (**Figures 5-6**). Among these changes, we identified a significant increase in LacdiNAcs and GlcNAcs with a loss of oncofetal epitope 3’sulfo-Lewis^A/C^. Conserved changes in glycosylation underlie many of the processes throughout cancer progression and can drive changes in cell phenotype through regulation of cell surface receptors^66^. Receptors that stimulate cell proliferation, growth, and oncogenesis have been shown to have more canonical N-glycosylation motif sites (N-X-S/T), longer extracellular domains, and an increased number of sites per 100 amino acids than other classes of receptor (e.g. those involved in vascular formation and organogenesis). N-glycan branching and multiplicity is highly dependent on metabolic flux via the hexosamine pathway^67^. A recent study demonstrated that induction of GNAS^R201C^ increases glycolysis in IPMN cell lines^68^. Modulating glucose metabolism could be one way that oncogenic GNAS modulates N-glycan composition.

N-glycosylation has been shown to differentially regulate EGFR/PI3K/ERK and TGF-β signaling, integrin-matrix interactions, pathways involved in cell proliferation, and EMT^67, 69^. Consistent with previous reports, we show here that mutant GNAS^R201C^ induces pyloric-like differentiation, inhibits cancer cell invasion, and drives an MET-like phenotype^14, 68, 70^. Concomitant with MET, we have identified significant shifts in N-glycosylation including a significant increase in LacdiNAc glycans. Cleavage of LacdiNAcs by chitinase results in a reversal of the MET phenotype suggesting that LacdiNAcs modify cell surface receptors involved in EMT and motility, perhaps by inhibiting activity^60^. Further studies are required to identify which receptors undergo LacdiNAc N-glycosylation and how this impacts downstream signaling.

We found that GNAS^R201C^ induces *B4Galnt3*, which promotes the formation of terminal LacdiNAcs, which prevents the synthesis of Lewis epitopes. This is accomplished by the addition of N-acetyl-galactose in lieu of galactose, required for the formation of Lewis epitopes. Recently, it has been shown that a capping domain on B4GALNT3 are important for shunting glycans towards LacdiNAc instead of Lewis epitopes^55^. Here, we found that these LacdiNacs led to a benign phenotype and that their removal with chitinase promoted the development of aggressive tumor behavior. In addition, others have demonstrated that forced expression of Lewis epitopes in murine PDAC models let to greater inflammation as well as oncogenic transformation, suggesting that Lewis epitopes are carcinogens^57^. Thus, it appears that GNAS induces an indolent phenotype in the KRAS background by (1) favoring the production of LacdiNAcs which promote a benign phenotype and (2) decreasing expression of acidic Lewis epitopes that promote inflammation and cancer. As such, we propose that GNAS acts as a glycan rheostat controlling oncogenic potential. To our knowledge this could be the first example of a genetic mutation that inhibits tumorigenesis by controlling glycan species abundance. While further studies are required to validate this proposed mechanism, we show here that LacdiNAcs (WFA) and 3’-sulfo-Le^A/C^ (Das-1) staining are non-overlapping and specifically label low-and high-grade IPMN, respectfully. Utilizing this combination of biomarkers in the future may aid in the risk stratification of IPMN patients for surgery; getting immediate care to those who need it while sparing others unnecessary surgery.

Altogether, this study provides insight into the distinct clinical behavior, cystic form, and mucin-rich nature of some IPMN. Mutant GNAS drives an indolent, pyloric phenotype in IPMN through a SPDEF-CREB3L1 axis with potentially anti-tumorigenic glycosylation changes. Further studies will evaluate cell surface receptor modification by glycans, how this impacts receptor activity, and if glycosylation can serve as a therapeutic target in addition to a biomarker.

## ABBREVIATIONS

ADM: acinar to ductal metaplasia
DOX: doxycycline
EEC: enteroendocrine cells
EMT: epithelial to mesenchymal transition
GSEA: gene set enrichment analysis
GEMMs: genetically engineered mouse models
GSII: Griffonia simplicifolia
HG: high grade
IPMN: intraductal papillary mucinous neoplasm
LG: low grade
MET: mesenchymal to epithelial transition
mxIHC: multiplex immunohistochemistry
PDAC: pancreatic ductal adenocarcinoma
PanIN: pancreatic intraepithelial neoplasia
ROI: region of interest
scRNA-seq: single cell RNA sequencing
SPEM: spasmolytic polypeptide expressing metaplasia
WFA: Wisteria floribunda

## ACKNOWLEDGEMENTS

The authors would like to thank Sydney Batts, JoAnna Dennis, Sergey Ivanov, and Nidhi Jyotsana for technical assistance. The Vanderbilt Creative Data Solutions Shared Resource (RRID:SCR_022366) performed and/or assisted with RNA-Seq data processing, analysis, and deposition to GEO. The Vanderbilt University Medical Center VANTAGE Core provided technical assistance for sample preparation and sequencing, and VANTAGE is supported in part by Clinical and Translational Science Award Grant 5UL1 RR024975-03, Vanderbilt Ingram Cancer Center Grant P30 CA68485, Vanderbilt Vision Center Grant P30 EY08126, and National Institutes of Health/National Center for Research Resources Grant G20 RR030956. Glycomics analysis was performed at the Complex Carbohydrate Research Center and was supported in part by the National Institutes of Health (NIH)-funded R24 grant (R24GM137782) to Parastoo Azadi.

## DATA AVAILABILITY STATEMENT

Sequencing data generated in this study is available in the gene expression omnibus (GSE244280). https://www.ncbi.nlm.nih.gov/geo/query/acc.cgi

## BACKGROUND AND CONTEXT

Intraductal papillary mucinous neoplasms (IPMN) are cystic precancerous lesions characterized by GNAS mutations. Markers that distinguish lesions that will progress to pancreatic cancer versus those that will remain benign are lacking.

## NEW FINDINGS

Mutant GNAS drives a mucinous pyloric phenotype in IPMN through SPDEF/CREB3L1 characterized by distinct changes in N glycosylation. GNAS^R201C^ expression upregulates LacdiNAc glycans at the expense of pro-tumorigenic Lewis antigens, driving an indolent phenotype.

## LIMITATIONS

Further studies are required to determine what cell surface receptors are impacted by GNAS^R201C^-driven N glycosylation and how this impacts downstream signaling.

## CLINICAL RESEARCH RELEVANCE

LacdiNAc and 3’-Sulfo-Le^A/C^ glycan expression are mutually exclusive in IPMN and distinguish low-from high-grade lesions. Combined use of these markers may aid in the risk stratification of patients to identify those requiring immediate treatment and surgical resection versus those that should be monitored.

## BASIC RESEARCH RELEVANCE

GNAS induces an indolent phenotype in IPMN by driving LacdiNAc glycan synthesis which promotes a benign phenotype and decreasing expression of acidic Lewis epitopes that promote inflammation and cancer. As such, we propose that GNAS acts as a glycan rheostat controlling oncogenic potential.

## Lay summary

This study provides insight into the distinct clinical behavior, cystic form, and mucin-rich nature of IPMN. Mutant GNAS drives anti-tumorigenic glycosylation which may be used to distinguish benign from malignant disease.

---

**Figure S1. Expression of pyloric metaplasia markers in human IPMN.** (**A**) Uniform manifold approximation and projection (UMAP) of single cell RNA-sequencing (scRNA-seq) data from 6 patients with IPMN and/or PDAC from Bernard et al. (**B**) Expression of epithelial marker *EPCAM* and (**C**) immune marker *PTPRC* (CD45). (**D**) Expression of mucins *MUC5AC*, *MUC6* (gastric folveolar IPMN), and *MUC2* (intestinal IPMN). (**E**) Expression of SPEM markers *TFF2*, *AQP5*, and *CD44*.

**Figure S2. Expression of pyloric metaplasia markers in human pre-malignant lesions and PDAC.** Heat map showing expression of SPEM markers identified in a murine model of pancreatitis (Ma et al.^20^) in a previously reported dataset of laser capture dissected epithelium from patient IPMN (n = 19), PanIN (n = 26), and PDAC (n = 197)^37^.

**Figure S3. Pyloric metaplasia markers are expressed in human IPMN.** (**A**) Representative images from hematoxylin, MUC5AC, CD44v9, and AQP5 from mxIHC separated by low grade gastric foveolar IPMN or high grade pancreatobiliary IPMN. Quantification of staining in (A) and Figure 1 plotted as (**B**) low grade vs. high grade or (**C**) by molecular subtype (gastric foveolar, intestinal, or pancreatobiliary). *, p < 0.05; ***, p < 0.005; ****, p < 0.001.

**Figure S4. GNAS^R201C^ expression drives transcriptomic changes in PDAC cell lines.** (**A**) Heat map of differentially expressed genes between cell lines 4838 and C241. (**B**) Expression of epithelial markers or (**C**) mesenchymal markers in 4838 or C241 cells +/- DOX treatment determined by RNA sequencing. (**D**) Expression of epithelial markers or (**E**) mesenchymal markers in 4838 or C241 cells +/- DOX treatment determined by qRT-PCR. (**F**) GSEA analysis of gene expression signature changes in 4838 or (**G**) C241 cells with DOX treatment. CTRL, control; DOX, doxycycline. *, p < 0.05; **, p < 0.01; ***, p < 0.005; **** p < 0.001.

**Figure S5. GNAS^R201C^ expression drives a mucinous, cystic phenotype.** (**A**) Quantification of highest-grade lesion or cystic score in *KrasGNAS* mice +/-DOX (combined 10- and 20-weeks treatment). Grade, 1 = 1/3 or tissue is low grade; 2 = 2/3 of tissue is low grade; 3 = high grade; 4 = invasive. Cystic score, 0 = no cysts; 1 = mile; 2 = extensive. (**B**) Heat map showing expression of select tuft cells, enteroendocrine cell (EEC), or mucus cell gene markers from Maurer et al.^37^, from the epithelium of human IPMN (n = 19), PanIN (n = 26), or PDAC (n = 197). (**C**) *MUC5AC*, *MUC5B*, and *MUC13* expression in PDAC tumors from cBioPortal with wild type (n = 96) or mutant GNAS^R201C/H^ (n = 4). *, p < 0.05; ***, p < 0.005; ****, p < 0.001.

**Figure S6. Creb3l1 and Creb3l4 are predicted regulators of SPEM.** RNA expression or binarized activity of (**A**) *Creb3l1* or (**B**) *Creb3l4* predicted by regulon analysis overlayed on the UMAP from Figure 3A.

**Figure S7. SPEM markers are predicted targets of Creb3l1 and Creb3l4.** Plots of either (**A**) *Creb3l1* or (**B**) *Creb3l4* predicted target genes generated from regulon analysis of scRNA-seq data generated form murine pancreatitis^20^. Select markers are circled in red.

**Figure S8. Spdef and Creb3l4 target genes increase in expression with GNAS^R201C^.** Heat maps of (**A**) *Spdef* target genes overlaid on RNA-seq data of 4838 (top) or C241 (bottom) cells +/- DOX and GNAS^R201C^ expression or (**B**) *Creb3l4* target genes overlaid on RNA-seq data of either 4838 (left) or C241 (right) cells +/- DOX. Top 10% of gene targets are shown. Select SPEM markers are bolded.

**Figure S9. Creb3l1 target genes increase in expression with GNAS^R201C^.** Heat maps of *Creb3l1* target genes overlaid on the RNA-seq data of either 4838 (left) or C241 (right) cells +/- DOX and GNAS^R201C^ expression. Top 10% of gene targets are shown. SPEM markers are highlighted.

**Figure S10. GNAS^R201C^ drives glycosyltransferase gene expression changes.** (**A**) Heat map with hierarchical clustering of glycosyltransferase gene expression in either 4838 or C241 cells treated with control or doxycycline (DOX). (**B**) Principal component analysis of RNA-seq generated from organoids from *Ptf1aCre/+;Rosa26R-LSL-rtTA-TetO-GNAS^R201C^* (*GNAS*) mice treated with either control or DOX *in vitro* (n = 2/condition). (**C**) Volcano plot of differentially expressed genes between control and DOX treated organoids. (**D**) Heat map of glycosyltransferase gene expression in *GNAS* organoids treated with either control or DOX. (**E**) qRT-PCR for human *GNAS* or LacdiNAc transferase enzyme *B4galnt3* in *GNAS* organoids treated with either control or DOX. *, p < 0.05; **, p < 0.01.

**Figure S11. GNAS^R201C^ expression drives minor O glycan changes.** (**A**) Principal component analysis of total O glycan species changes between control and DOX treated 4838 or C241 cells. (**B**) Representative spectra of O glycosylation mass spectrometry in C241 cells; control, top and DOX-treated, bottom. (**C**) O glycans and modifications identified in 4838 cells +/- DOX. (**D**) O glycans and modifications identified in C241 cells +/- DOX. *, p < 0.05; **, p < 0.01.

**Figure S12. GNAS^R201C^ expression drives major N glycan changes.** (**A**) N glycan modifications identified in 4838 cells +/- DOX. (**B**) Representative spectra of N glycosylation mass spectrometry in C241 cells; control, left and DOX-treated, right. (**C**) N glycan modifications identified in C241 cells +/- DOX. *, p < 0.05.

**Figure S13. PanIN and IPMN formation are accompanied by significant changes in N glycan deposition.** (**A**) Hematoxylin and eosin (H&E) staining of pancreata from *KrasGNAS* mice on either control or DOX chow for 22 weeks. Scale bars, 100 μm. (**B**) Spectra collected by imaging N glycosylation mass spectrometry of acinar enriched areas or PanIN from *KrasGNAS* mice on control chow or IPMN enriched areas from *KrasGNAS* mice on DOX chow. (**C**) Quantification of predicted glycans characterized by fucosylation or (**D**) additional glycans. (**E**) Representative heat map of expression of a predicted LacdiNAc (control tissues, top; DOX chow, bottom) and quantification of identified species. (**F**) Quantification of predicted glycan structures characterized by sialyation. *, p < 0.05; **, p < 0.01.

**Figure S14. Spdef, Creb3l1, and Chitinase impact cell phenotype and LacdiNAc abundance.** (**A**) qRT-PCR for human *GNAS* in either 4838 or C241 cell treated with either control (CTRL) or doxycycline (DOX) or DOX+ control siRNA or DOX+ siRNAs against *Spdef* or *Creb3l1*. (**B**) qRT-PCR for *Epcam*, *Cldn2*, or *Zeb1* in 4838 or C241 cells treated with either DOX + control siRNA or siRNAs against *Spdef* or (**C**) *Creb3l1*. (**D**) Quantification of lectin WFA (LacdiNAcs) area and signal intensity in 4838 or (**E**) C241 cells treated with DOX and either control or *Spdef* siRNA. (**F**) Quantification of WFA area and signal intensity in 4838 or (**G**) C241 cells treated with DOX and either control or *Creb3l1* siRNA. *, p < 0.05; **, p < 0.01; ***, p < 0.005; ****, p < 0.001.

**Table S1.** Primary antibodies used for western blot (WB), immunohistochemistry (IHC), or immunofluorescence (IF) studies.

| **Antibody** | **Company** | **Catalog #** | **Dilution,**<br>**WB** | **Dilution,**<br>**IHC or IF** |
| --- | --- | --- | --- | --- |
| AQP5 | Sigma-Aldrich | HPA065008 | N/A | 1:500 |
| CD44v9 (human) | Cosmo Bio | LKG-M003 | N/A | 1:250 |
| CD44v10e16 (mouse) | Cosmo Bio | LKG-M002 | N/A | 1:250 |
| CREB3L1 | Sigma | HPA024069 | N/A | 1:300 |
| CREB3L4 | Sigma | HPA038122 | N/A | 1:300 |
| Das-1 | Leinco | N/A | N/A | 1:500 |
| DCLK1 | Abcam | Ab37994 | N/A | 1:500 |
| GAPDH | Cell Signaling | 5174 | 1:1000 | N/A |
| GNAS (R201C) | GeneTex | GTX135412 | 1:1000 | N/A |
| GSII | Invitrogen | L21415 | N/A | 1:250 |
| KRAS (G12D) | Cell Signaling | 14429 | 1:1000 | N/A |
| MUC5AC | Invitrogen | MA-12178 | N/A | 1:500 |
| SPDEF | Lifespan Bioscience | LS-C-499857 | N/A | 1:300 |
| Synaptophysin (SYP) | Cell Marque | 336R-94 | N/A | 1:500 |
| WFA | Invitrogen | L32481 | N/A | 1:250 |

**Table S2.**
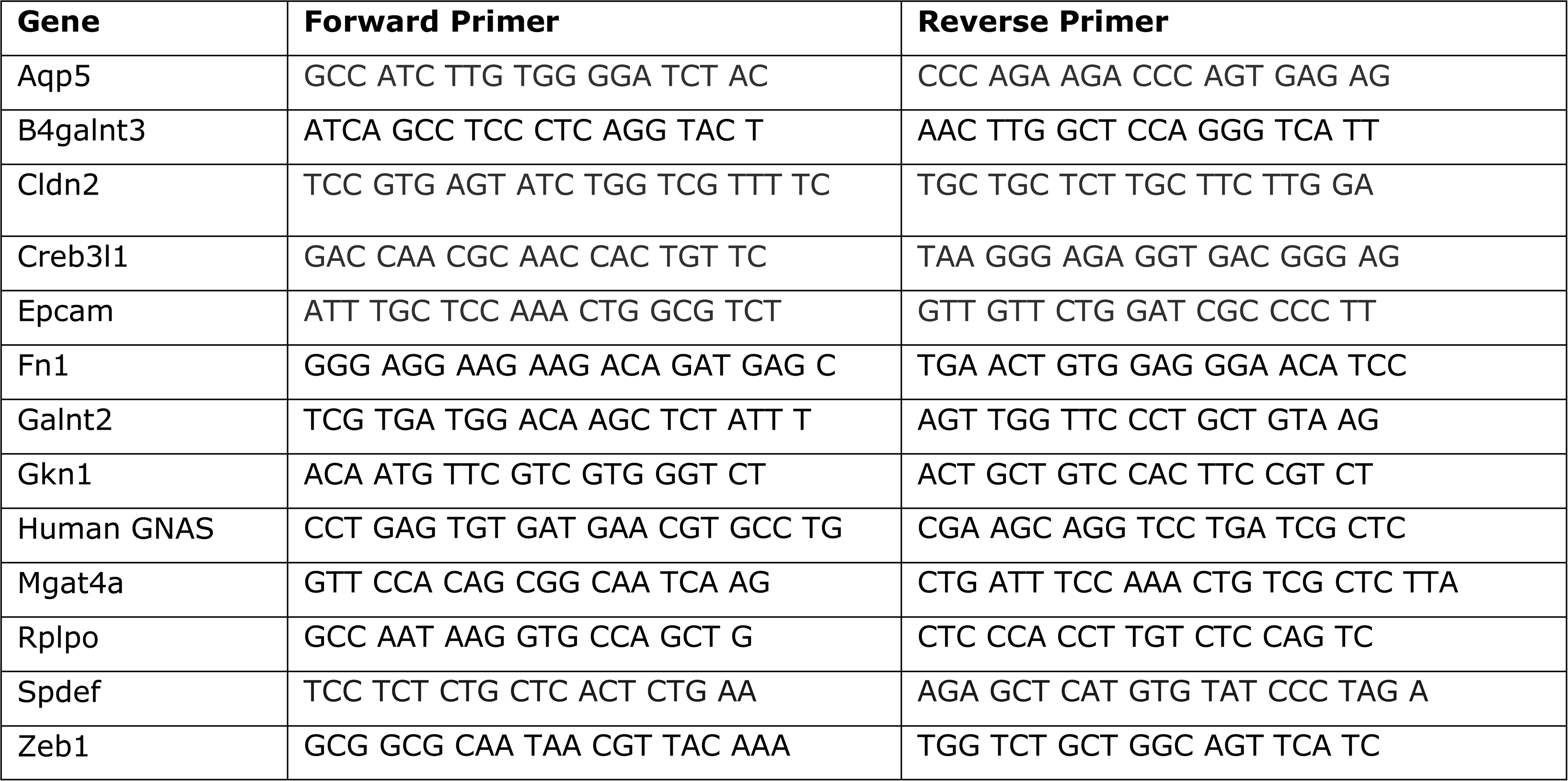
Primers for qRT-PCR.

